# Optimisation of GMP-compliant hollow-fiber culturing and scaled-up separation of mesenchymal stem cell-derived extracellular vesicles to overcome unexpected impact of bovine serum contaminants′ transfer to end products

**DOI:** 10.64898/2026.11.15.766728

**Authors:** Alan Rivera-Tenorio, Ainara González-Moro, Miriam M. Rodríguez de Lope, Eduardo Chicano-Gálvez, Javier Martin Ramirez, Jose Carlos Cabrera, Fernando de la Cuesta

## Abstract

The clinical translation of mesenchymal stem cell-derived extracellular vesicles (MSC-EVs) requires scalable manufacturing strategies ensuring product purity, reproducibility, and compliance with Good Manufacturing Practice (GMP) standards. However, fetal bovine serum (FBS)-derived contaminants may persist in EV preparations, compromising their characterization and clinical applicability. This study aimed to optimize MSC-EV production using hollow-fiber bioreactors while minimizing bovine protein contamination through improved culture and purification strategies. Extracellular vesicles from immortalized human adipose-derived MSCs were produced under different culture conditions, including FBS supplementation, partial replacement with Panexin, and use of chemically defined medium for high-density cultures (CDM-HD). EVs were purified using tangential flow filtration, fast protein liquid chromatography, and ultrafiltration, and characterized by nanoparticle tracking analysis, electron microscopy, Western blot, proteomics, and ELISA.

Proteomic analysis revealed substantial bovine protein contamination under conventional FBS-containing conditions, with 11.4% of identified proteins exclusively assigned to bovine origin. Transitioning to CDM-HD reduced this proportion to 2.6%, markedly decreasing albumin abundance. Optimization of Panexin supplementation supported MSC expansion while reducing the necessary percentage of bovine serum. Although albumin became undetectable by Western blot, sensitive ELISA measurements revealed residual contamination and suggested unexpected protein carryover associated with chromatography column reuse. Importantly, using a new purification column yielded CDM-HD-derived EV preparations with albumin concentrations below ELISA assay’s quantification limit (0.69 ng/mL). These findings highlight the importance of controlling both upstream culture conditions and downstream purification to prevent exogenous protein contamination, establishing a promising scalable manufacturing workflow for high-purity MSC-EVs suitable for further GMP-oriented development and future clinical translation.

## INTRODUCTION

Mesenchymal stem cells (MSCs) have attracted considerable interest in regenerative medicine due to their remarkable regenerative, immunomodulatory, and anti-inflammatory properties, making them promising candidates for the treatment of a broad spectrum of diseases (Harrell et al. 2020; Gowen et al. 2020). Although the therapeutic potential of MSCs was initially attributed to their capacity to engraft into injured tissues and differentiate into specialized cell types, accumulating evidence has demonstrated that their beneficial effects are primarily mediated through paracrine mechanisms rather than direct cell replacement (Allan et al. 2020; Giebel and Lim 2025). Among the components of the MSC secretome, extracellular vesicles (EVs) have emerged as the principal mediators of this intercellular communication, transporting proteins, lipids, messenger RNAs, microRNAs, and other bioactive molecules capable of modulating the phenotype and function of recipient cells (Welsh et al. 2024; Allan et al. 2020). Consequently, MSC-derived EVs (MSC-EVs) have gained increasing attention as a cell-free therapeutic platform capable of reproducing many of the beneficial biological activities of their parental cells while potentially overcoming several limitations associated with cell-based therapies, including lower immunogenicity, improved biosafety, and easier handling and storage (Gowen et al. 2020; Massa et al. 2020; Giebel and Lim 2025).

The growing interest in MSC-EVs has accelerated their translation from experimental research toward clinical application, leading to a rapid increase in preclinical studies and early-phase clinical trials investigating their therapeutic potential across a wide range of pathological conditions (Giebel and Lim 2025; Shimizu et al. 2026). A screening of ClinicalTrials.gov records conducted on October 30, 2026, identified 28 interventional clinical studies investigating MSC-derived EVs. These studies explore their therapeutic potential in a broad range of conditions, including respiratory diseases (acute respiratory distress syndrome, COVID-19, bronchopulmonary dysplasia and pulmonary fibrosis), liver failure, inflammatory bowel diseases (Crohn’s disease and ulcerative colitis), neurological disorders (ischemic stroke and neurodegenerative diseases), dermatological conditions (burns, dystrophic epidermolysis bullosa and psoriasis), as well as ocular, auditory, periodontal and metabolic disorders (ClinicalTrials.gov, accessed October 30, 2026). However, the successful clinical implementation of MSC-EVs requires the development of manufacturing processes capable of consistently producing vesicle preparations that are safe, reproducible, scalable, and compliant with Good Manufacturing Practice (GMP) standards (Rohde et al. 2019; Giebel and Lim 2025). Unlike conventional laboratory-scale production, clinical-grade manufacturing demands strict control over every step of the process, including cell expansion, culture conditions, EV isolation, purification, characterization, storage, and quality control, all of which directly influence the composition, functionality, and therapeutic efficacy of the final product (Shimizu et al. 2026; Cecchin et al. 2023). Consequently, optimizing EV manufacturing has become one of the major challenges limiting the widespread clinical translation of MSC-EV-based therapies and has stimulated the development of new culture systems and production strategies aimed at improving both yield and product quality (Rohde et al. 2019; Giebel and Lim 2025).

The clinical translation of MSC-derived EVs requires manufacturing processes capable of generating products with consistent quality, safety, and biological activity while complying with regulatory standards for advanced biological therapeutics (Adlerz et al. 2020; Wiest and Zubair 2020). Achieving this objective requires careful control of every stage of the production process, including cell sourcing, expansion conditions, culture medium composition, EV isolation, downstream purification, storage, and quality control, as each of these variables can directly influence the physicochemical characteristics, molecular composition, and therapeutic performance of the final EV preparation (Shimizu et al. 2026; Chua et al. 2022). Consequently, considerable efforts have focused on developing standardized and scalable manufacturing strategies that ensure batch-to-batch reproducibility while meeting Good Manufacturing Practice (GMP) requirements for future clinical application (Costa-Ferro et al. 2024; Shimizu et al. 2026).

Among the different variables involved in MSC-EV manufacturing, cell culture conditions have received particular attention because they directly influence both EV production and product quality (Shimizu et al. 2026; Wiest and Zubair 2020). Fetal bovine serum (FBS) remains one of the most widely used supplements for MSC expansion due to its ability to provide essential nutrients and growth factors that support cell proliferation and viability (Eitan et al. 2015). However, FBS also contains abundant bovine proteins and extracellular vesicles that cannot be completely removed by conventional EV-depletion protocols, allowing these components to co-isolate with cell-derived EVs and interfere with downstream molecular characterization and functional analyses (Lehrich et al. 2018; Driedonks et al. 2019; Pham et al. 2021a). The persistence of serum-derived contaminants has therefore become an important concern for the manufacture of clinical-grade MSC-EVs because it compromises product purity, complicates the interpretation of experimental results, and represents an additional challenge for standardization and regulatory compliance (Pham et al. 2021a; Wiest and Zubair 2020; Chua et al. 2022).

To address the limitations associated with FBS supplementation, increasing efforts have focused on the development of serum-free and chemically defined culture media for MSC expansion and EV production (Yoon et al. 2025; Figueroa-Valdés et al. 2021). Unlike conventional serum-containing media, chemically defined formulations eliminate animal-derived components, thereby reducing batch-to-batch variability and minimizing the introduction of exogenous contaminants while facilitating compliance with Good Manufacturing Practice (GMP) requirements for clinical manufacturing (Yoon et al. 2025; Calascibetta et al. 2025). Several studies have demonstrated that MSCs cultured under chemically defined conditions maintain their biological properties while supporting EV production and biological activity, although differences in the molecular composition of isolated EV preparations have been reported depending on the culture conditions employed (Figueroa-Valdés et al. 2021; Zhu et al. 2021a; Calascibetta et al. 2025). Consequently, chemically defined media have emerged as a promising strategy to improve the standardization and clinical translation of MSC-EV manufacturing (Yoon et al. 2025; Calascibetta et al. 2025). In this regard, FiberCell Systems recognizes bovine serum albumin (BSA) as a potential contaminant associated with serum-containing culture media and recommends CDM-HD, a protein-free medium, as a replacement for FBS in hollow fiber bioreactors to minimize serum-derived protein contamination (FiberCell Systems, 2018). However, the extent to which these culture conditions reduce residual serum-derived contaminants within isolated EV preparations remains to be fully established (Yoon et al. 2025; Calascibetta et al. 2025).

Although chemically defined media address an important source of variability during MSC-EV production, their implementation alone does not overcome another major challenge associated with clinical translation: the development of scalable manufacturing processes. Conventional two-dimensional flask-based culture systems remain the most widely used platform for MSC expansion; however, they are labor-intensive, require extensive culture surface area, and are difficult to standardize for the large-scale production of extracellular vesicles (Wiest and Zubair 2020; Jakl et al. 2023). To address these limitations, hollow fiber bioreactors have emerged as a promising manufacturing platform by providing a high-surface-area three-dimensional culture environment that supports high-density cell culture and enables the continuous harvesting of conditioned medium containing cell-secreted EVs for subsequent downstream isolation and purification, thereby improving production efficiency while reducing manual handling and facilitating process standardization(Garcia et al. 2024; Gobin et al. 2021; Jakl et al. 2023). Consequently, the integration of chemically defined culture media with hollow fiber bioreactor technology represents a promising strategy for establishing scalable and GMP-compatible manufacturing workflows for MSC-EVs. However, the impact of combining these manufacturing strategies on the purity of isolated MSC-EV preparations remains insufficiently characterized, particularly with respect to the persistence of residual serum-derived protein contamination (Garcia et al. 2024; Wiest and Zubair 2020).

Despite the considerable advances achieved in both culture media optimization and scalable manufacturing technologies, important challenges remain to produce clinical-grade MSC-EVs. Although chemically defined media and hollow fiber bioreactor systems have independently emerged as promising strategies to improve manufacturing processes, the impact of different upstream manufacturing workflows on the purity of MSC-EV preparations remains poorly characterized, particularly regarding residual serum-derived contaminants. Addressing this question is essential for the development of standardized manufacturing workflows capable of generating EV preparations that meet the quality requirements expected for future GMP production and clinical translation.

Therefore, the aim of this study was to compare two distinct manufacturing workflows to produce MSC-derived extracellular vesicles using the same hollow fiber bioreactor platform. These workflows differed in their upstream culture strategies, including cell adaptation procedures, extracellular matrix coating, and culture medium composition, while maintaining an identical downstream EV isolation process. MSC-EV preparations were comparatively characterized by nanoparticle tracking analysis, soluble protein quantification, Western blot analysis of EV and contaminant markers, and quantitative proteomic profiling to determine how different manufacturing workflows influence EV purity and residual serum-derived protein contamination under scalable production conditions.

## MATERIALS AND METHODS

### 2.1. Cell Culture

Immortalized human adipose tissue-derived mesenchymal stem cells (AT-MSCs; ATCC) were initially expanded according to the manufacturer’s instructions using Mesenchymal STEM Basal Medium supplemented with the Mesenchymal Stem Cell Growth Kit for Adipose and Umbilical-derived MSCs–Low Serum (ATCC). This medium contained recombinant human basic FGF (5 ng/mL), acidic FGF (5 ng/mL), EGF (5 ng/mL), 2% fetal bovine serum (FBS), and L-alanyl-L-glutamine (2.4 mM). Cells were maintained at 37°C in a humidified atmosphere containing 5% CO₂, and culture medium was replaced every 48 h.After recovery and initial expansion, the commercial medium was completely replaced with laboratory-prepared DMEM (4.5 g/L glucose, Corning) supplemented with 2% FBS (Gibco), 5 ng/mL EGF (PeproTech), 5 ng/mL acidic FGF (PeproTech), 5 ng/mL basic FGF (Sigma-Aldrich), 100 U/mL penicillin (Corning), 100 μg/mL streptomycin (Corning), and 2.5 μg/mL amphotericin B (Sigma-Aldrich.

Cells were subsequently divided into two experimental groups. One population was expanded in DMEM supplemented with 10% FBS for inoculation into the first hollow-fiber bioreactor 20 ml of high flux Polysulfone cartridge (Fiber Cell). The second population remained in DMEM containing 2% FBS and was subjected to a gradual adaptation to Panexin-containing medium before inoculation into the second hollow-fiber bioreactor cartridge.

### 2.2. Adaptation to a reduced-serum chemically defined medium

Cells assigned to the second hollow-fiber bioreactor were progressively adapted from DMEM supplemented with 2% FBS and growth factors to media containing increasing concentrations of Panexin (PanBiotech) and decreasing concentrations of FBS. Adaptation was performed over four sequential passages, maintaining the cells in each medium composition for three passages before proceeding to the next condition, by gradually replacing FBS with Panexin as follows: 1.5% FBS + 0.5% Panexin // 1.0% FBS + 1.0% Panexin // 0.5% FBS + 1.5% Panexin // 0.2% FBS + 1.8% Panexin.

To facilitate the adaptation process, two stock media were prepared. The first consisted of DMEM supplemented with 2% FBS, 5 ng/mL EGF, 5 ng/mL acidic FGF, 5 ng/mL basic FGF, 100 U/mL penicillin, 100 μg/mL streptomycin and 2.5 μg/mL amphotericin B. The second consisted of DMEM supplemented with 2% Panexin, 100 U/mL penicillin, 100 μg/mL streptomycin and 2.5 μg/mL amphotericin B. Appropriate volumes of each stock medium were mixed to obtain the desired FBS/Panexin ratios.

### 2.3 Assessment of cell viability during medium adaptation

Cell viability under each adaptation condition was subsequently assessed by Crystal Violet staining. AT-MSCs were seeded in duplicate 24-well plates at a density of 8,000 cells per well in complete culture medium containing 2% FBS. After 8 h, the medium was removed, cells were washed once with PBS, and the culture medium was replaced with one of the four adaptation conditions (1.5% FBS/0.5% Panexin, 1.0% FBS/1.0% Panexin, 0.5% FBS/1.5% Panexin, or 0.2% FBS/1.8% Panexin). One plate was processed immediately after medium replacement (day 0), while the second plate was cultured for 5 days under the corresponding adaptation conditions with medium renewal every 48 h before analysis (day 5). At each time point, relative cell density was determined by Crystal Violet staining.

Following the incubation period, the culture medium was removed, and the cells were washed once with PBS before fixation with 500 μL of 1% glutaraldehyde (VWR Chemicals) for 10 min at room temperature. Cells were then washed three times with PBS and stained with 500 μL of 1% Crystal Violet (Merck) for 30 min at room temperature. Excess dye was removed by washing the wells with distilled water, and the plates were allowed to air dry.

The retained dye was solubilized with 2 mL of 10% acetic acid (PanReac AppliChem) for 5 min. After homogenization, 250 μL of each sample were transferred in technical duplicate to a 96-well plate, using 10% acetic acid as blank control test sample. Absorbance was measured at 595 nm using a FLUOstar OPTIMA microplate reader (BMG LABTECH).

### 2.4 Hollow-fiber bioreactor inoculation

Two 20 mL high flux Polysulfone cartridges (FiberCell Systems) were used in a hollow-fiber bioreactor system to evaluate two different culture conditions in separate experiments.

#### First hollow-fiber bioreactor (10% FBS condition)

The first hollow-fiber bioreactor was used for AT-MSCs cultured in DMEM supplemented with 10% FBS. Before cell inoculation, the cartridge was prepared following a three-day conditioning protocol. On the first day, both the intracapillary space (ICS) and extracapillary space (ECS) were filled with sterile PBS and maintained for 24 h. The PBS used to fill the ECS was supplemented with 1 mg fibronectin to promote cell attachment. After 24 h, PBS was replaced in both compartments with serum-free DMEM supplemented only with penicillin (100 U/mL), streptomycin (100 µg/mL) and amphotericin B (2.5 µg/mL), and the bioreactor was incubated for an additional 24 h. On the third day, both compartments were filled with the corresponding complete culture medium and maintained for a further 24 h before cell inoculation.

AT-MSCs expanded in DMEM supplemented with 10% FBS were cultured in T175 tissue culture flasks (STL) until reaching the required cell number for inoculation. A total of 5 × 10⁷ cells were harvested and inoculated into the ECS compartment of the bioreactor. Approximately 1 h after inoculation, continuous medium perfusion was initiated by connecting the circulation pump. During the expansion phase, the central medium reservoir initially contained 200 mL of the corresponding culture medium, which was replaced every 48-h. Cell growth was monitored indirectly by measuring glucose consumption using a GlucCell™ glucose meter (Esco). As the metabolic activity of the culture increased, the daily perfusion volume was progressively adjusted according to glucose consumption. Whenever glucose concentration decreased to approximately 50% of its initial value, the circulating medium volume was increased to ensure an adequate nutrient supply. This process was repeated until a maximum perfusion volume of 1 L/day was reached.

#### Second hollow-fiber bioreactor (Panexin condition)

Following the initial experiment using 10% FBS, a second hollow-fiber bioreactor was used to evaluate AT-MSCs adapted to DMEM supplemented with 0.5% FBS and 1.5% Panexin. The cartridge was prepared following the same three-day conditioning protocol described above, with the exception that the PBS used to fill the ECS was not supplemented with fibronectin.

AT-MSCs expanded in DMEM supplemented with 0.5% FBS and 1.5% Panexin were cultured in T175 tissue culture flasks (STL) until reaching the required cell number for inoculation. A total of 5 × 10⁷ cells were harvested and inoculated into the ECS compartment of the bioreactor. Approximately 1 h after inoculation, continuous medium perfusion was initiated by connecting the circulation pump. During the expansion phase, the central medium reservoir initially contained 200 mL of the corresponding culture medium, which was replaced every 48 h. Cell growth was monitored indirectly by measuring glucose consumption using a GlucCell™ glucose meter (Esco). As the metabolic activity of the culture increased, the daily perfusion volume was progressively adjusted according to glucose consumption. Whenever glucose concentration decreased to approximately 50% of its initial value, the circulating medium volume was increased to ensure an adequate nutrient supply. This process was repeated until a maximum perfusion volume of 1 L/day was reached.

Following cell expansion, cultures were gradually adapted to Chemically Defined Medium–High Density (CDM-HD, FiberCell Systems) to eliminate serum-derived contaminants prior to extracellular vesicle (EV) production. Medium adaptation was performed stepwise by sequentially replacing the original culture medium with increasing proportions of CDM-HD (25%, 50%, 75%, and finally 100%). Progression between adaptation steps was determined by the metabolic performance of the culture, assessed through daily glucose monitoring. Once complete adaptation to CDM-HD was achieved, the bioreactors were maintained under continuous perfusion with 1 L/day of fresh CDM-HD for EV production.

### 2.5 Extracellular Vesicle Isolation

EV-enriched conditioned medium was harvested daily from the extracapillary space (ECS) of the hollow-fiber bioreactors starting 7 days after cell inoculation. To recover the conditioned medium retained within the ECS, 20 mL of serum-free DMEM was introduced into the compartment, allowing displacement of the EV-enriched conditioned medium, which was subsequently collected. Samples were centrifuged at 2,000 × g for 10 min at 4°C to remove cell debris, and the resulting supernatants were stored at −80°C until EV isolation. Before purification, frozen conditioned media were thawed at room temperature and filtered through a 0.45 μm membrane filter. The samples were subsequently concentrated using a TFF-Easy 15 kDa tangential flow filtration system (HansaBioMed) until the desired volume was reached. Purification of EVs was performed by fast protein liquid chromatography (FPLC) using an ÄKTA Start system (Cytiva) equipped with a Capto Core 700 column (Cytiva). Prior to each run, the column was equilibrated with PBS, and concentrated samples were manually loaded using a 5 mL injection loop. Elution was carried out using PBS as the mobile phase, and 1.5 mL fractions were collected throughout the chromatographic run. During the initial purification runs, protein concentration in the collected fractions was assessed using the Pierce BCA Protein Assay Kit (Thermo Fisher Scientific) to verify the chromatographic fractionation profile and identify fractions with protein concentrations below 25 µg/µL. Once the correspondence between the chromatographic profile and the fractions suitable for collection had been established, BCA measurements were discontinued, and fraction selection in subsequent runs was based solely on the chromatographic profile. Fractions corresponding to the EV-containing peak were subsequently concentrated using Amicon Ultra 100 KDa centrifugal filters (Sigma-Aldrich) until a final volume of approximately 1–1.5 mL was obtained. Concentrated EV preparations were then stored at −80°C until further characterization and downstream experiments.

### 2.6 EV Characterization

EV preparations were characterized in accordance with the Minimal Information for Studies of Extracellular Vesicles (MISEV2023) guidelines by means of nanoparticle tracking analysis (NTA), transmission electron microscopy (TEM) and Western blot analysis of EV identity markers.

#### 2.6.1 Nanoparticle tracking analysis (NTA)

Particle concentration and size distribution were determined by NTA using a NanoSight LM10 instrument equipped with a NanoSight Syringe Pump (Malvern Panalytical). EV samples were diluted in PBS (typically 1:20–1:50) to obtain the optimal particle concentration for analysis, and five 60-s videos were acquired under continuous flow. Particle concentration (particles/mL), mean particle diameter, and mode size were determined using NanoSight software.

#### 2.6.2 Transmission electron microscopy (TEM)

EV morphology was evaluated by transmission electron microscopy (TEM). Samples were fixed with 0.1% paraformaldehyde, deposited onto Formvar/carbon-coated copper grids (Ted Pella), negatively stained with 2% uranyl acetate, and visualized using a JEM-1010 transmission electron microscope (JEOL).

#### 2.6.3 Western blot of EV identity markers and bovine serum albumin

EV samples were lysed by adding 1% SDS (Sigma-Aldrich) and 1% protease inhibitor cocktail (Sigma-Aldrich) to the final concentrated EV suspension (50 μL). Samples were incubated on ice for 5 min and subsequently sonicated using a Fisherbrand Ultrasonic Bath Select 30 (Fisher Scientific) in three cycles of 2 min, with 2 min on ice between each cycle. Lysates were centrifuged at 12,000 rpm for 5 min at 4°C, and the supernatants were recovered. The total protein concentration of EV preparations was determined using the Pierce BCA Protein Assay Kit (Thermo Fisher Scientific) according to the manufacturer’s instructions, using bovine serum albumin (BSA) as the protein standard.

To confirm EV identity, equal amounts of protein (40 μg) were separated on 10% SDS-PAGE gels under non-reducing conditions and transferred onto PVDF membranes. Membranes were blocked with 5% BSA, except for BSA detection, for which 5% milk was used as the blocking solution. Membranes were then incubated overnight at 4°C with primary antibodies against the EV markers CD63 (Abcam, 1:500), CD81 (Santa Cruz Biotechnology, 1:100), calnexin (Santa Cruz Biotechnology, 1:5000) and BSA (Invitrogen, 1:1000). After incubation with the corresponding HRP-conjugated secondary antibodies, protein bands were detected using enhanced chemiluminescence (ECL, NZYtech).

### 2.8 Proteomic analyses

EV protein extracts were obtained as described above for Western blot analysis. Protein concentration was determined using the Qubit™ Protein Assay Kit (Thermo Fisher Scientific). Protein digestion was performed using the iST Sample Preparation Kit (PreOmics) according to the manufacturer’s instructions. The resulting peptides were resuspended in 0.1% formic acid to a final concentration of 10 ng/µL, and 400 ng of peptides were loaded onto Evotips (Evosep Biosystems). Peptide separation was performed using an Evosep One liquid chromatography system (Evosep Biosystems) equipped with a PepSep analytical column (150 µm × 150 mm, 1.5 µm particle size) maintained at 60°C and operated with the 88 samples per day gradient. The LC system was coupled online to a timsTOF fleX mass spectrometer (Bruker Daltonics) operated in data-independent acquisition parallel accumulation–serial fragmentation (DIA-PASEF) mode. Instrument performance was monitored using HeLa Protein Digest Standard (Evosep Biosystems) and iRT peptides (Biognosys).

Proteomic analyses were performed at the Proteomics Unit from IMIBIC, Córdoba, using independent EV preparations obtained under each experimental condition. Each biological replicate was analyzed in technical triplicates by LC–MS/MS. Protein identification and label-free quantification were performed using Spectronaut™ 18 (Biognosys) with the directDIA workflow against the UniProt human protein database. Carbamidomethylation of cysteine residues was set as a fixed modification, whereas methionine oxidation and protein N-terminal acetylation were considered variable modifications. Protein and peptide identifications were filtered using a false discovery rate (FDR) of 1%.

### 2.9 ELISA albumin quantification essay

Residual BSA in the EV preparations was quantified using a commercially available BSA ELISA kit (Bethyl Laboratories, Inc.), according to the manufacturer’s instructions. EVs produced under the different culture conditions were analyzed, including EVs generated in medium supplemented with 1.5% Panexin and 0.5% FBS and EVs produced under CDM 10% conditions. A BSA sample diluted 1:250,000 was included as a positive control to verify the functionality of the assay. The BSA standard was reconstituted at 500 ng/mL and serially diluted 1:3 to generate standards ranging from 500 to 0.69 ng/mL, with dilution buffer serving as the blank. Samples, standards, and the positive control were analyzed in duplicate. Appropriately diluted samples and standards (100 μL/well) were incubated for 1 h at room temperature (20–25 °C), followed by four washes with 1× wash buffer. Detection antibody (100 μL/well) was added and incubated for 1 h at room temperature, followed by four washes. Subsequently, HRP Solution C (100 μL/well) was added and incubated for 30 min at room temperature, followed by four additional washes. TMB substrate (100 μL/well) was then added and the plate was incubated for 30 min at room temperature in the dark. The reaction was stopped by adding 100 μL/well of stop solution, and absorbance was measured at 450 nm using a BioTek Epoch 2 microplate reader (BioTek Instruments, Inc.) within 30 min of stopping the reaction. Albumin concentrations were determined by interpolation from a four-parameter logistic (4PL) standard curve and corrected for the corresponding sample dilution factor. Only values falling within the assay’s standard-curve range (0.69–500 ng/mL) were considered quantitatively reliable. Samples with signals below the lowest standard (0.69 ng/mL) were considered below the quantifiable range of the assay and were not extrapolated to obtain numerical concentrations.

### 2.10 Statistical analyses

Statistical analyses were performed using GraphPad Prism 8 (GraphPad Software), except for the proteomic data, which were analyzed using Spectronaut 18 and MetaboAnalyst. Data are expressed as mean ± standard error of the mean (SEM), with n representing the number of independent experiments conducted in each case. Normality of data distribution was assessed using the Kolmogorov–Smirnov test for comparisons between two groups and the Shapiro–Wilk test for one-way ANOVA. Homogeneity of variances was assessed using the Brown–Forsythe test. Comparisons between two groups were performed using an unpaired two-tailed Student’s *t*-test. For comparisons involving more than two groups, one-way analysis of variance (ANOVA) followed by Dunnett’s multiple comparisons test was used. Differences were considered statistically significant at *p* < 0.05.

## RESULTS

### 1. AT-MSCs derived EVs isolation and characterization

Following concentration of the conditioned medium collected from the hollow-fiber bioreactor, EVs were isolated by FPLC using a Capto Core 700 column (Figure 1A). The isolated EV preparations were subsequently characterized by nanoparticle tracking analysis (NTA), transmission electron microscopy (TEM), and Western blot.

**Figure 1.**
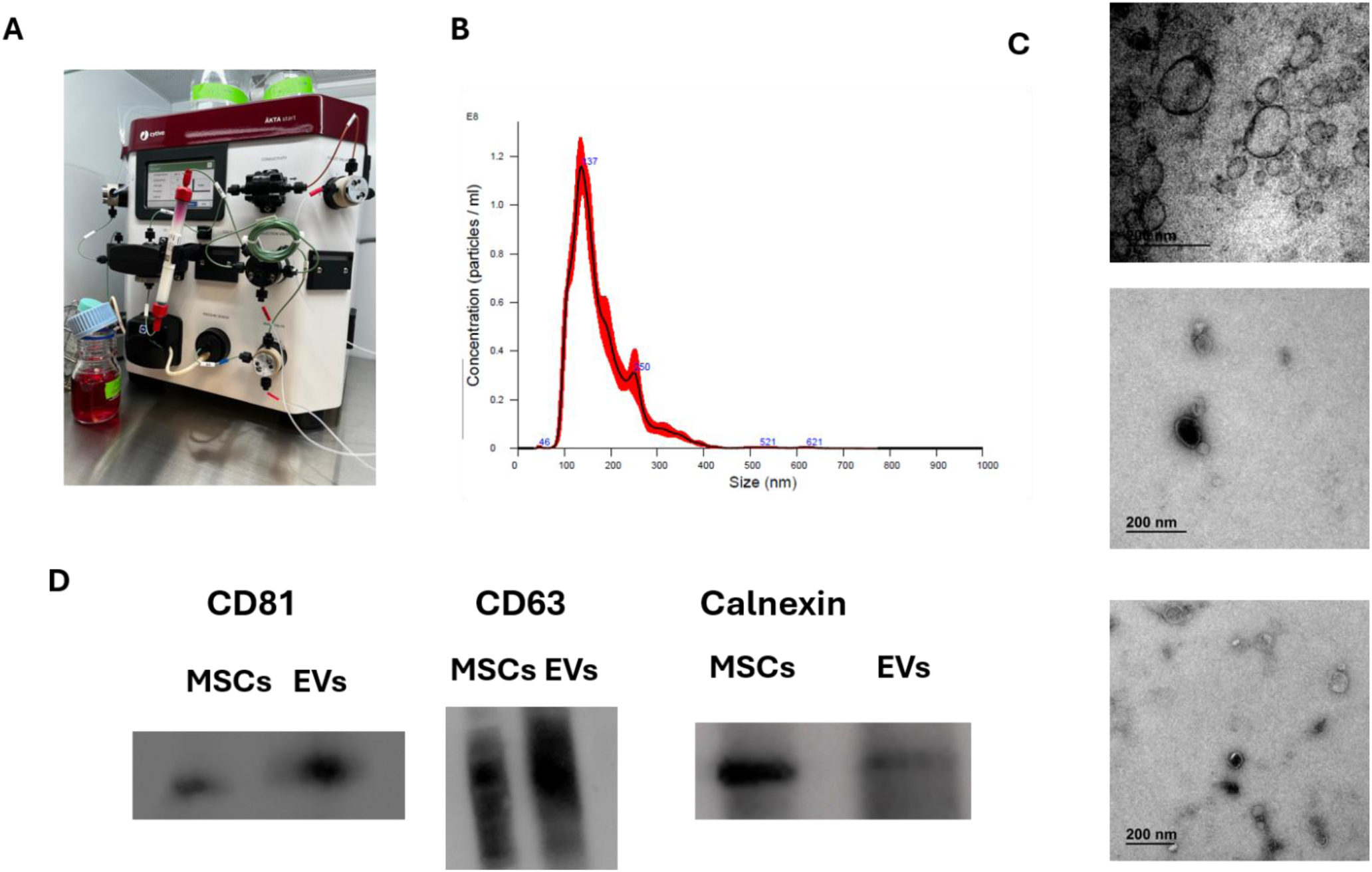
Isolation and characterization of AT-MSC-derived EVs from hollow-fiber bioreactor conditioned medium. (A) Representative image of the concentrated conditioned medium during FPLC fractionation using a Capto Core 700 column. (B) Representative NTA profile showing the particle concentration and size distribution of EVs isolated from conditioned medium containing 10% CDM-HD. (C) Representative TEM image of EVs isolated from conditioned medium containing 10% CDM-HD. (D) Representative Western blot showing the EV-associated markers CD81 and CD63 and the cellular marker calnexin in EVs isolated from 10% CDM-HD-conditioned medium (right lane) and AT-MSC cell lysate (left lane).

NTA analysis confirmed the presence of a nanoparticle population with a modal size of approximately 169 nm and a concentration of 1.5 x 10^10^ particles/mL (Figure 1B). TEM analysis further confirmed the presence of nanosized vesicular structures consistent with EV morphology (Figure 1C).

The presence of EV-associated proteins was further assessed by Western blot. The EV-associated markers CD81 and CD63 were detected in the EV preparation obtained from conditioned medium containing 10% CDM-HD, whereas the cellular marker calnexin was detected in the AT-MSC cell lysate but was barely detected in the EV preparation (Figure 1D). Together, these results support the successful isolation and characterization of AT-MSC-derived EVs from hollow-fiber bioreactor conditioned medium.

### 2. Proteomic characterization of MSC-EVs produced with 10% FBS circulating medium shows presence of bovine proteins

To characterize the protein composition of EVs produced under standard culture conditions, EVs isolated from conditioned medium collected from the hollow-fiber bioreactor maintained in 10% FBS were subjected to LC-MS/MS-based proteomic analysis. A total of 809 proteins were identified in the isolated EV preparation. Of these, 448 proteins (55.4%) were unambiguously assigned to *Homo sapiens*, whereas 92 proteins (11.4%) were exclusively assigned to *Bos taurus* (Figure 3A). The remaining 269 proteins (33.3%) could not be unambiguously assigned to a single species, mainly due to shared or non-specific peptide sequences between human and bovine proteins. The detection of bovine proteins in the isolated EV preparation was unexpected given the use of a hollow-fiber bioreactor with a nominal 20-kDa molecular weight cut-off and prompted further investigation of potential serum-derived protein carryover.

Among the 35 most abundant proteins identified in the dataset, several proteins associated with extracellular matrix and EV preparations were detected together with multiple bovine proteins (Table 1). Notably, BSA was the fourth most abundant protein identified in the dataset, making it the most highly represented bovine protein among the top 35 identifications. Other bovine proteins detected among these highly abundant proteins included alpha-2-HS-glycoprotein, alpha-1-antiproteinase, alpha-fetoprotein, fibulin-1, and tenascin-C. In contrast, several of the most abundant human proteins corresponded to extracellular matrix proteins, including fibronectin, decorin, collagens, laminins, and tenascin-C, as well as proteins previously reported in EV proteomes. The presence of abundant bovine serum-derived proteins, particularly albumin, indicated that components originating from the FBS were present in the isolated material under these culture conditions and motivated subsequent experiments aimed at evaluating and reducing serum-derived protein contamination.

**Table 1.** Most abundant proteins identified by LC-MS/MS proteomic analysis of EVs produced in the hollow-fiber bioreactor under 10% FBS culture conditions. The 35 proteins with the highest abundance in the proteomic dataset are shown. Protein identity and organism assignment were based on UniProt annotations. EV relevance was classified according to reported evidence of association with EV preparations, extracellular matrix localization, or potential FBS-derived origin. Color coding indicates species assignment: green, proteins unambiguously assigned to *Homo sapiens*; red, proteins unambiguously assigned to *Bos taurus*; yellow, proteins for which species assignment was ambiguous due to shared or non-specific peptide sequences between human and bovine proteins.

| Ranking | UniProt ID | Protein name | Organism | EV relevance |
|---|---|---|---|---|
| 1 | P13611 | Versican core protein | *Homo sapiens* | EV-associated / ECM |
| 2 | P02751 | Fibronectin | *Homo sapiens* | EV-associated / ECM |
| 3 | P07585 | Decorin | *Homo sapiens* | EV-associated / ECM |
| 4 | A0A140T897;P02769 | Albumin | *Bos taurus* | Potential FBS-derived serum protein |
| 5 | P12763 | Alpha-2-HS-glycoprotein | *Bos taurus* | Potential FBS-derived serum protein |
| 6 | P02452 | Collagen alpha-1(I) chain | *Homo sapiens* | EV-associated / ECM |
| 7 | Q08380 | Galectin-3-binding protein | *Homo sapiens* | EV-associated / ECM |
| 8 | P24821 | Tenascin | *Homo sapiens* | EV-associated / ECM |
| 9 | P23142 | Fibulin-1 | *Homo sapiens* | EV-associated / ECM |
| 10 | P04264 | Keratin, type II cytoskeletal 1 | *Homo sapiens* | EV-associated / cytoskeletal; non-specific |
| 11 | Q7Z7G0 | Target of Nesh-SH3 (extracellular) | *Homo sapiens* | Extracellular/ECM; EV relevance uncertain |
| 12 | P13645 | Keratin, type I cytoskeletal 10 | *Homo sapiens* | EV-associated / cytoskeletal; non-specific |
| 13 | P60709;P60712;P63258;P63261 | Actin, cytoplasmic 1/2 | *Homo sapiens; Bos taurus* | Mixed-species / cytoskeletal; non-specific |
| 14 | P34955 | Alpha-1-antiproteinase | *Bos taurus* | Potential FBS-derived serum protein |
| 15 | Q3SZ57 | Alpha-fetoprotein | *Bos taurus* | Potential FBS-derived serum protein |
| 16 | P11047 | Laminin subunit gamma-1 | *Homo sapiens* | EV-associated / ECM |
| 17 | P08962 | CD63 antigen | *Homo sapiens* | Strong EV-associated protein |
| 18 | P98160 | Basement membrane-specific heparan sulfate proteoglycan core protein | *Homo sapiens* | EV-associated / ECM |
| 19 | P07477;P07478;Q8NHM4 | Serine protease 1; Trypsin-2; Putative trypsin-6 | *Homo sapiens* | Potential technical/biological contaminant |
| 20 | Q562R1 | Beta-actin-like protein 2 | *Homo sapiens* | Cellular/cytoskeletal; non-specific |
| 21 | A0A3Q1M4K3;P0CG47;P0CG48;P0CG53;P0CH28;P62979;P62987;P62992;P63048 | Ubiquitin / polyubiquitin / ubiquitin-ribosomal fusions | *Bos taurus; Homo sapiens* | Mixed-species / cellular; non-specific |
| 22 | P07942 | Laminin subunit beta-1 | *Homo sapiens* | EV-associated / ECM |
| 23 | P08123 | Collagen alpha-2(I) chain | *Homo sapiens* | EV-associated / ECM |
| 24 | P35527 | Keratin, type I cytoskeletal 9 | *Homo sapiens* | EV-associated / cytoskeletal; non-specific |
| 25 | A0A3Q1MJB2 | Fibulin-1 | *Bos taurus* | Potential FBS-derived extracellular/ECM protein |
| 26 | P12109 | Collagen alpha-1(VI) chain | *Homo sapiens* | EV-associated / ECM |
| 27 | P51884 | Lumican | *Homo sapiens* | EV-associated / ECM |
| 28 | A0A3Q1LKE5 | Tenascin C | *Bos taurus* | Potential FBS-derived extracellular/ECM protein |
| 29 | O00468 | Agrin | *Homo sapiens* | EV-associated / ECM |
| 30 | P02461 | Collagen alpha-1(III) chain | *Homo sapiens* | EV-associated / ECM |
| 31 | P01024 | Complement C3 | *Homo sapiens* | EV-associated / complement-associated; non-specific |
| 32 | P02533 | Keratin, type I cytoskeletal 14 | *Homo sapiens* | EV-associated / cytoskeletal; non-specific |
| 33 | Q2TBN8 | E3 ubiquitin-protein ligase FANCL | *Bos taurus* | Cellular protein; EV relevance uncertain |
| 34 | A5D7R6 | ITIH2 protein | *Bos taurus* | Potential FBS-derived serum/extracellular protein |
| 35 | P12111 | Collagen alpha-3(VI) chain | *Homo sapiens* | EV-associated / ECM |

### 3. Impairment of identity marker detection by bovine proteins on MSC-EVs derived from cells cultured in 10% FBS circulating medium, in comparison to 10% CDM-HD

To evaluate the removal of FBS-derived contaminants during the transition to chemically defined medium (CDM-HD), EV preparations collected at different stages of medium replacement were analyzed by Western blot. The EV markers CD63 and CD81 were evaluated in EV preparations to confirm the presence of extracellular vesicles, whereas bovine serum albumin (BSA), the major protein component of fetal bovine serum (FBS), was evaluated as an indicator of residual serum contamination.

As shown in Figure 2, albumin was detected in EV preparations isolated while the bioreactor was maintained in 10% FBS (lane 1). Albumin signal was still detected in EVs collected during the transition to 50% FBS/50% CDM-HD (lanes 2 and 3), although with lower intensity. In contrast, no albumin signal was detected in EV preparations obtained after complete adaptation of the bioreactor to 10% CDM-HD (lanes 4 and 5), indicating effective removal of FBS-derived protein contaminants.

**Figure 2.**
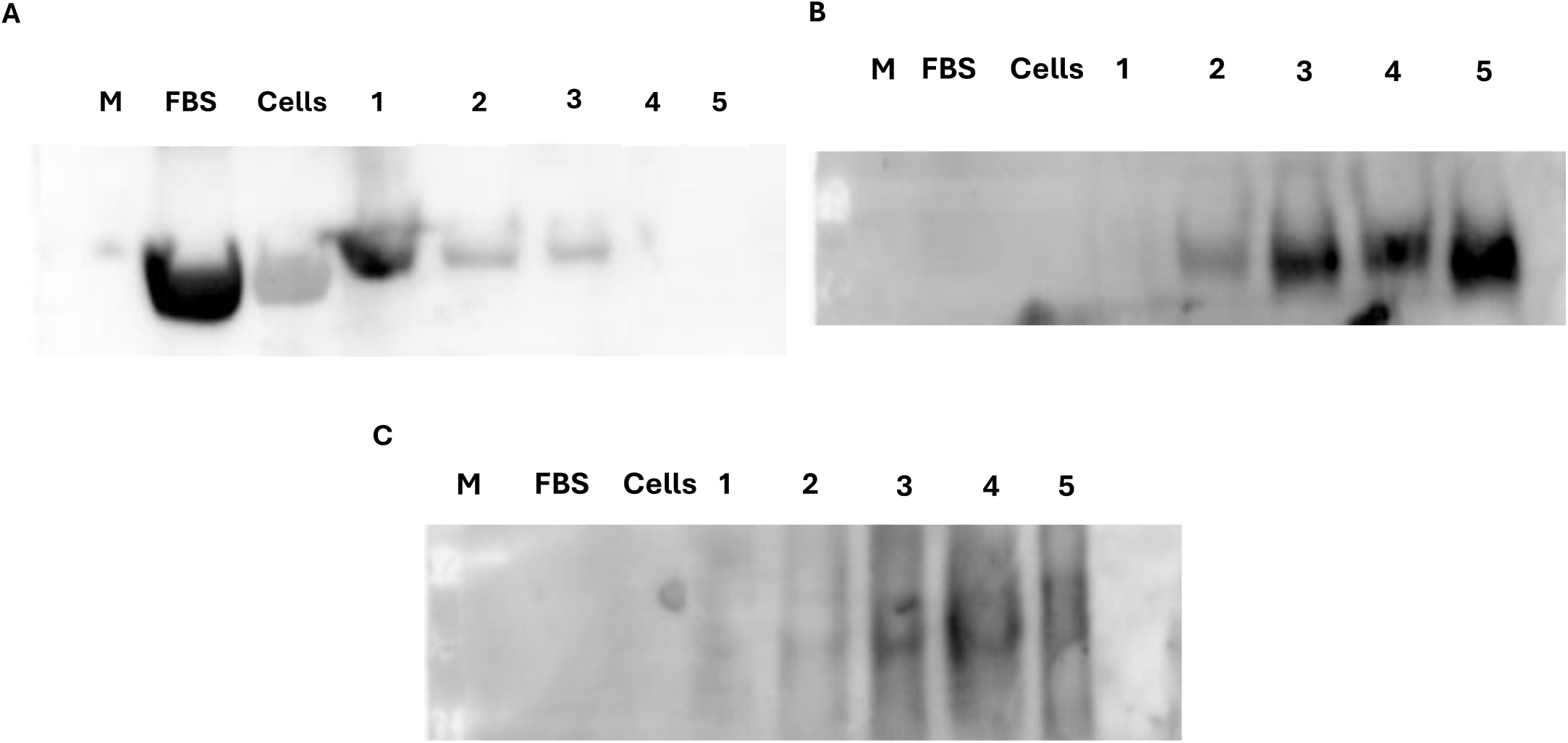
Western blot analysis of albumin and EV markers in EVs collected during the transition from 10% FBS-containing medium to 10% CDM-HD. (A) Albumin immunoblot. (B) CD81 immunoblot. (C) CD63 immunoblot. Lane M, molecular weight marker; lane FBS, fetal bovine serum; lane Cells, AT-MSC cell lysate; lane 1, EVs isolated from cultures maintained in 10% FBS; lanes 2–3, EVs isolated during the transition to CDM-HD (5% FBS/5% CDM-HD); lanes 4–5, EVs isolated after complete adaptation to 10% CDM-HD.

**Figure 3.**
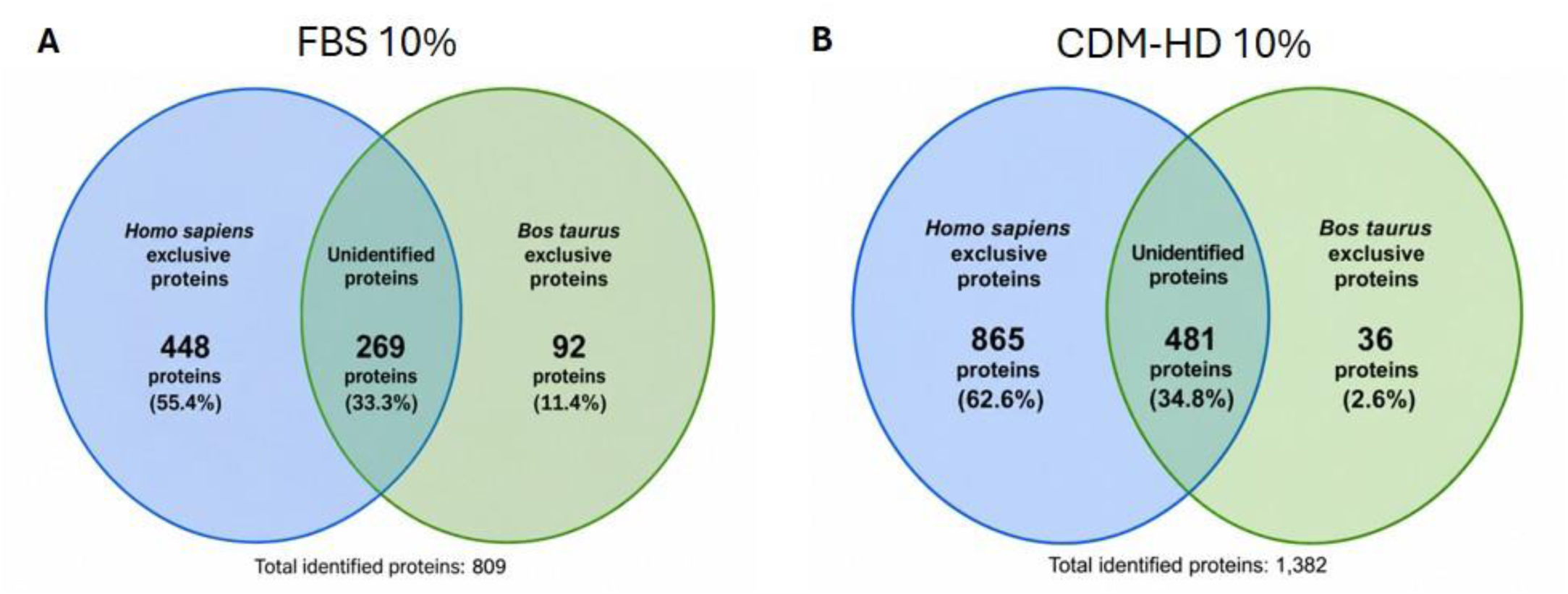
Comparative analysis of the protein distribution identified by proteomic analysis of extracellular vesicles (EVs) produced under different culture conditions. (A) EVs obtained in medium supplemented with 10% FBS, with 809 proteins identified. (B) EVs obtained in medium supplemented with 10% CDM-HD, with 1,382 proteins identified. Venn diagrams show the distribution of proteins classified as Homo sapiens-specific, Bos taurus-specific, or unidentified, expressed as the number of proteins and the percentage of the total identified proteins under each condition.

The EV markers CD81 and CD63 were detected in all EV preparations. CD81 was detected in all EV preparations, with stronger signal intensity observed in EVs isolated under CDM-HD conditions than in FBS-containing samples. CD63 was detected as a broad band between approximately 37 and 75 kDa in all EV preparations, consistent with its characteristic migration pattern and confirming the successful isolation of EVs regardless of the culture medium.

### 4. Proteomic analysis of EVs derived from cells cultured in 10% CDM-HD circulating medium, after initial expansion in 10% FBS

To evaluate the protein composition of EVs following the transition from FBS-containing medium to chemically defined medium, EVs isolated from conditioned medium collected from the hollow-fiber bioreactor maintained in 10% CDM-HD were subjected to LC-MS/MS-based proteomic analysis. A total of 1,382 proteins were identified in the isolated material. Of these, 865 proteins (62.6%) were unambiguously assigned to *Homo sapiens*, whereas only 36 proteins (2.6%) were unambiguously assigned to *Bos taurus*. The remaining 481 proteins (34.8%) could not be unambiguously assigned to either species based on the identified peptide sequences (Figure 3B).

Compared with the proteomic profile obtained under 10% FBS conditions, the number and proportion of unambiguously identified bovine proteins were markedly reduced following the transition to CDM-HD. In the previous 10% FBS condition, 92 proteins were assigned to *Bos taurus* (11.4% of the identified proteins), whereas only 36 bovine proteins (2.6%) were detected under 10% CDM-HD conditions. Conversely, the number of unambiguously assigned human proteins increased from 448 (55.4%) to 865 (62.6%). Although the proportion of ambiguous identifications remained substantial, the reduction in bovine assignments suggests a lower contribution of serum-derived proteins to the isolated material under CDM-HD conditions.

Analysis of the 35 most abundant protein identifications revealed a predominance of human proteins associated with extracellular matrix and cellular components (Table 2). Notably, bovine Tenascin C (A0A3Q1LKE5) was ranked fifth among all protein identifications, making it the only bovine protein detected within the 35 most abundant proteins. In contrast, BSA (A0A140T897; P02769), which represented the fourth most abundant protein under the previous 10% FBS condition, was ranked 485th under 10% CDM-HD conditions. This marked decrease in the relative abundance of albumin is consistent with a substantial reduction in serum-derived protein contribution following the transition to chemically defined medium.

**Table 2.** Most abundant proteins identified in EVs produced under 10% CDM-HD culture conditions. The 35 proteins with the highest abundance in the proteomic dataset are shown. Protein identity and organism assignment were based on UniProt annotations. EV relevance was classified according to reported evidence of association with EV preparations, extracellular matrix localization, or potential FBS-derived origin. Color coding: green, proteins unambiguously assigned to *Homo sapiens*; red, proteins unambiguously assigned to *Bos taurus*; yellow, proteins for which species assignment was ambiguous due to shared or non-specific peptide sequences between human and bovine proteins.

| Ranking | UniProt ID | Protein name | Organism | EV relevance |
|---|---|---|---|---|
| 1 | P24821 | Tenascin | *Homo sapiens* | EV-associated / ECM |
| 2 | Q08380 | Galectin-3-binding protein | *Homo sapiens* | EV-associated / ECM |
| 3 | P02794 | Ferritin heavy chain | *Homo sapiens* | Cellular/serum-associated; non-specific |
| 4 | P02751 | Fibronectin | *Homo sapiens* | EV-associated / ECM |
| 5 | A0A3Q1LKE5 | Tenascin C | *Bos taurus* | Potential FBS-derived extracellular/ECM protein |
| 6 | P02792 | Ferritin light chain | *Homo sapiens* | Cellular/serum-associated; non-specific |
| 7 | P12109 | Collagen alpha-1(VI) chain | *Homo sapiens* | EV-associated / ECM |
| 8 | P13611 | Versican core protein | *Homo sapiens* | EV-associated / ECM |
| 9 | P00338 | L-lactate dehydrogenase A chain | *Homo sapiens* | EV-associated / metabolic; non-specific |
| 10 | P04406 | Glyceraldehyde-3-phosphate dehydrogenase | *Homo sapiens* | EV-associated / metabolic; non-specific |
| 11 | P12814 | Alpha-actinin-1 | *Homo sapiens* | EV-associated / cytoskeletal; non-specific |
| 12 | P11047 | Laminin subunit gamma-1 | *Homo sapiens* | EV-associated / ECM |
| 13 | P16070 | CD44 antigen | *Homo sapiens* | EV-associated / membrane; non-specific |
| 14 | P07195 | L-lactate dehydrogenase B chain | *Homo sapiens* | EV-associated / metabolic; non-specific |
| 15 | P04075 | Fructose-bisphosphate aldolase A | *Homo sapiens* | EV-associated / metabolic; non-specific |
| 16 | O43707 | Alpha-actinin-4 | *Homo sapiens* | EV-associated / cytoskeletal; non-specific |
| 17 | Q562R1 | Beta-actin-like protein 2 | *Homo sapiens* | Cellular/cytoskeletal; non-specific |
| 18 | P60709;P60712;P63258;P63261 | Actin, cytoplasmic 1;Actin, cytoplasmic 1;Actin, cytoplasmic 2;Actin, cytoplasmic 2 | *Homo sapiens; Bos taurus* | Mixed-species / cytoskeletal; non-specific |
| 19 | P07942 | Laminin subunit beta-1 | *Homo sapiens* | EV-associated / ECM |
| 20 | O14818 | Proteasome subunit alpha type-7 | *Homo sapiens* | Cellular/proteasomal; non-specific |
| 21 | Q16363 | Laminin subunit alpha-4 | *Homo sapiens* | EV-associated / ECM |
| 22 | P09603 | Macrophage colony-stimulating factor 1 | *Homo sapiens* | Extracellular/cytokine; EV relevance uncertain |
| 23 | P12110 | Collagen alpha-2(VI) chain | *Homo sapiens* | EV-associated / ECM |
| 24 | Q99436 | Proteasome subunit beta type-7 | *Homo sapiens* | Cellular/proteasomal; non-specific |
| 25 | P12111 | Collagen alpha-3(VI) chain | *Homo sapiens* | EV-associated / ECM |
| 26 | P26022 | Pentraxin-related protein PTX3 | *Homo sapiens* | EV-associated / extracellular; non-specific |
| 27 | P98160 | Basement membrane-specific heparan sulfate proteoglycan core protein | *Homo sapiens* | EV-associated / ECM |
| 28 | P06733 | Alpha-enolase | *Homo sapiens* | EV-associated / metabolic; non-specific |
| 29 | O00468 | Agrin | *Homo sapiens* | EV-associated / ECM |
| 30 | P25789;Q3ZCK9 | Proteasome subunit alpha type-4 | *Homo sapiens; Bos taurus* | Mixed-species / proteasomal; non-specific |
| 31 | Q9Y6C2 | EMILIN-1 | *Homo sapiens* | EV-associated / ECM |
| 32 | P07585 | Decorin | *Homo sapiens* | EV-associated / ECM |
| 33 | P20618 | Proteasome subunit beta type-1 | *Homo sapiens* | Cellular/proteasomal; non-specific |
| 34 | A0A3Q1M4K3;P0CG47;P0CG48;P0CG53;P0CH28;P62979;P62987;P62992;P63048 | Ubiquitin-like domain-containing protein;Polyubiquitin-B;Polyubiquitin-C;Polyubiquitin-B;Polyubiquitin-C;Ubiquitin-40S ribosomal protein S27a;Ubiquitin-60S ribosomal protein L40;Ubiquitin-ribosomal protein eS31 fusion protein;Ubiquitin-ribosomal protein eL40 fusion protein | *Homo sapiens; Bos taurus* | Mixed-species / cellular; non-specific |
| 35 | P15144 | Aminopeptidase N | *Homo sapiens* | EV-associated / membrane/cellular; non-specific |

To further assess the changes in protein composition following the transition from FBS-containing to chemically defined medium, the 35 most abundant proteins identified under the 10% FBS condition were used as a reference, and their relative ranking was compared with that observed under 10% CDM-HD (Table 3). Overall, a marked redistribution of the most abundant proteins was observed following the change in culture conditions. Among the 35 proteins initially ranked highest under 10% FBS, 8 increased their relative ranking under 10% CDM-HD, 23 decreased in ranking, 3 were no longer detected, and 1 maintained the same position.

**Table 3.** Comparison of the 35 most abundant proteins identified under 10% FBS and 10% CDM-HD conditions. The 35 proteins with the highest ranking under the 10% FBS condition were used as the reference set, and their corresponding ranking under 10% CDM-HD was determined. Rank change represents the difference between the FBS and CDM-HD rankings; positive values indicate an increase in relative ranking (↑), whereas negative values indicate a decrease (↓). LOST indicates proteins identified among the top 35 under 10% FBS that were not detected in the 10% CDM-HD proteomic dataset. Species assignment was based on UniProt annotation, with green indicating *Homo sapiens*, red indicating *Bos taurus*, and yellow indicating ambiguous species assignment. Rankings represent relative positions within each individual proteomic dataset and should not be interpreted as direct comparisons of absolute protein abundance between conditions.

| **Ranking** | **Protein name** | **UniProt ID** | **Organism** | **CDM-HD rank** | **Rank change** | **Movement** | **Interpretation** |
|---|---|---|---|---|---|---|---|
| 1 | Versican core protein | P13611 | *Homo sapiens* | 8 | -7 | ↓ 7 | Moved down 7 positions |
| 2 | Fibronectin | P02751 | *Homo sapiens* | 4 | -2 | ↓ 2 | Moved down 2 positions |
| 3 | Decorin | P07585 | *Homo sapiens* | 32 | -29 | ↓ 29 | Moved down 29 positions |
| 4 | Albumin | A0A140T897;P02769 | *Bos taurus* | 485 | -481 | ↓ 481 | Moved down 481 positions |
| 5 | Alpha-2-HS-glycoprotein | P12763 | *Bos taurus* | 306 | -301 | ↓ 301 | Moved down 301 positions |
| 6 | Collagen alpha-1(I) chain | P02452 | *Homo sapiens* | 134 | -128 | ↓ 128 | Moved down 128 positions |
| 7 | Galectin-3-binding protein | Q08380 | *Homo sapiens* | 2 | 5 | ↑ 5 | Moved up 5 positions |
| 8 | Tenascin | P24821 | *Homo sapiens* | 1 | 7 | ↑ 7 | Moved up 7 positions |
| 9 | Fibulin-1 | P23142 | *Homo sapiens* | 171 | -162 | ↓ 162 | Moved down 162 positions |
| 10 | Keratin, type II cytoskeletal 1 | P04264 | *Homo sapiens* | 87 | -77 | ↓ 77 | Moved down 77 positions |
| 11 | Target of Nesh-SH3 (extracellular) | Q7Z7G0 | *Homo sapiens* | 51 | -40 | ↓ 40 | Moved down 40 positions |
| 12 | Keratin, type I cytoskeletal 10 | P13645 | *Homo sapiens* | 97 | -85 | ↓ 85 | Moved down 85 positions |
| 13 | Actin, cytoplasmic 1/2 | P60709;P60712;P63258;P63261 | *Homo sapiens; Bos taurus* | 18 | -5 | ↓ 5 | Moved down 5 positions |
| 14 | Alpha-1-antiproteina | P34955 | *Bos taurus* | 1012 | -998 | ↓ 998 | Moved down 998 positions |
| 15 | Alpha-fetoprotein | Q3SZ57 | *Bos taurus* |  |  | LOST | Not detected in CDM-HD |
| 16 | Laminin subunit gamma-1 | P11047 | *Homo sapiens* | 12 | 4 | ↑ 4 | Moved up 4 positions |
| 17 | CD63 antigen | P08962 | *Homo sapiens* | 39 | -22 | ↓ 22 | Moved down 22 positions |
| 18 | Basement membrane-specific heparan sulfate proteoglycan core protein | P98160 | *Homo sapiens* | 27 | -9 | ↓ 9 | Moved down 9 positions |
| 19 | Serine protease 1; Trypsin-2; Putative trypsin-6 | P07477;P07478;Q8NHM4 | *Homo sapiens* | 77 | -58 | ↓ 58 | Moved down 58 positions |
| 20 | Beta-actin-like protein 2 | Q562R1 | *Homo sapiens* | 17 | 3 | ↑ 3 | Moved up 3 positions |
| 21 | Ubiquitin / polyubiquitin / ubiquitin-ribosomal fusions | A0A3Q1M4K3;P0CG47;P0CG48;P0CG53;P0CH28;P62979;P62987;P62992;P63048 | *Homo sapiens; Bos taurus* | 34 | -13 | ↓ 13 | Moved down 13 positions |
| 22 | Laminin subunit beta-1 | P07942 | *Homo sapiens* | 19 | 3 | ↑ 3 | Moved up 3 positions |
| 23 | Collagen alpha-2(I) chain | P08123 | *Homo sapiens* | 179 | -156 | ↓ 156 | Moved down 156 positions |
| 24 | Keratin, type I cytoskeletal 9 | P35527 | *Homo sapiens* | 93 | -69 | ↓ 69 | Moved down 69 positions |
| 25 | Fibulin-1 | A0A3Q1MJB2 | *Bos taurus* |  |  | LOST | Not detected in CDM-HD |
| 26 | Collagen alpha-1(VI) chain | P12109 | *Homo sapiens* | 7 | 19 | ↑ 19 | Moved up 19 positions |
| 27 | Lumican | P51884 | *Homo sapiens* | 334 | -307 | ↓ 307 | Moved down 307 positions |
| 28 | Tenascin C | A0A3Q1LKE5 | *Bos taurus* | 5 | 23 | ↑ 23 | Moved up 23 positions |
| 29 | Agrin | O00468 | *Homo sapiens* | 29 | 0 | → 0 | No rank change |
| 30 | Collagen alpha-1(III) chain | P02461 | *Homo sapiens* | 166 | -136 | ↓ 136 | Moved down 136 positions |
| 31 | Complement C3 | P01024 | *Homo sapiens* | 113 | -82 | ↓ 82 | Moved down 82 positions |
| 32 | Keratin, type I cytoskeletal 14 | P02533 | *Homo sapiens* | 673 | -641 | ↓ 641 | Moved down 641 positions |
| 33 | E3 ubiquitin-protein ligase FANCL | Q2TBN8 | *Bos taurus* | 440 | -407 | ↓ 407 | Moved down 407 positions |
| 34 | ITIH2 protein | A5D7R6 | *Bos taurus* |  |  | LOST | Not detected in CDM-HD |
| 35 | Collagen alpha-3(VI) chain | P12111 | *Homo sapiens* | 25 | 10 | ↑ 10 | Moved up 10 positions |

The most pronounced decrease in relative ranking was observed for serum-derived proteins. BSA, which was the fourth most abundant protein under 10% FBS, decreased to rank 485 under 10% CDM-HD (↓481 positions). Similarly, bovine alpha-fetoprotein, fibulin-1, and ITIH2, which were ranked #15, #25, and #34 under 10% FBS, respectively, were no longer detected under CDM-HD conditions. All three proteins were assigned to *Bos taurus*, supporting a reduction in the contribution of bovine serum-derived proteins following the transition to chemically defined medium.

In contrast, several proteins associated with the extracellular matrix increased their relative ranking under CDM-HD. Tenascin, which ranked #8 under 10% FBS, became the highest-ranked protein in the CDM-HD dataset (#1), while galectin-3-binding protein increased from #7 to #2 and collagen alpha-1(VI) chain increased from #26 to #7. Notably, bovine tenascin C was not among the 35 most abundant proteins under the 10% FBS condition but ranked #5 under 10% CDM-HD, representing the only bovine protein among the 35 most abundant proteins in the CDM-HD dataset.

Together, these changes indicate a substantial shift in the relative composition of the most abundant proteins following replacement of FBS-containing medium with CDM-HD. The marked decrease in the relative ranking of albumin and the disappearance of several bovine serum-associated proteins, together with the increased representation of human extracellular matrix-associated proteins, suggest a reduced contribution of FBS-derived proteins to the EV preparation under CDM-HD conditions.

### 5. Optimization of chemically defined medium Panexin supplementation for AT-MSC culture

Aiming to minimize the observed bovine protein transference to the final MSC-EV end-product, we decided to investigate the use of a chemically defined medium, Panexin, that could replace FBS on the initial low-density stage of the hollow-fiber bioreactor culture. To identify a suitable Panexin supplementation strategy for AT-MSC culture, cells were progressively adapted to Panexin, maintaining them in medium containing increasing Panexin/FBS ratios for three passages before proceeding to the next condition. On each adaptation step, a subset of cells were maintained for 5 days in media containing a certain Panexin/FBS ratio. Representative phase-contrast images of each of these conditions, acquired on day 0 and day 5 are shown in Figure 4A, while cell biomass quantified by Crystal Violet staining is presented in Figure 4B.

**Figure 4.**
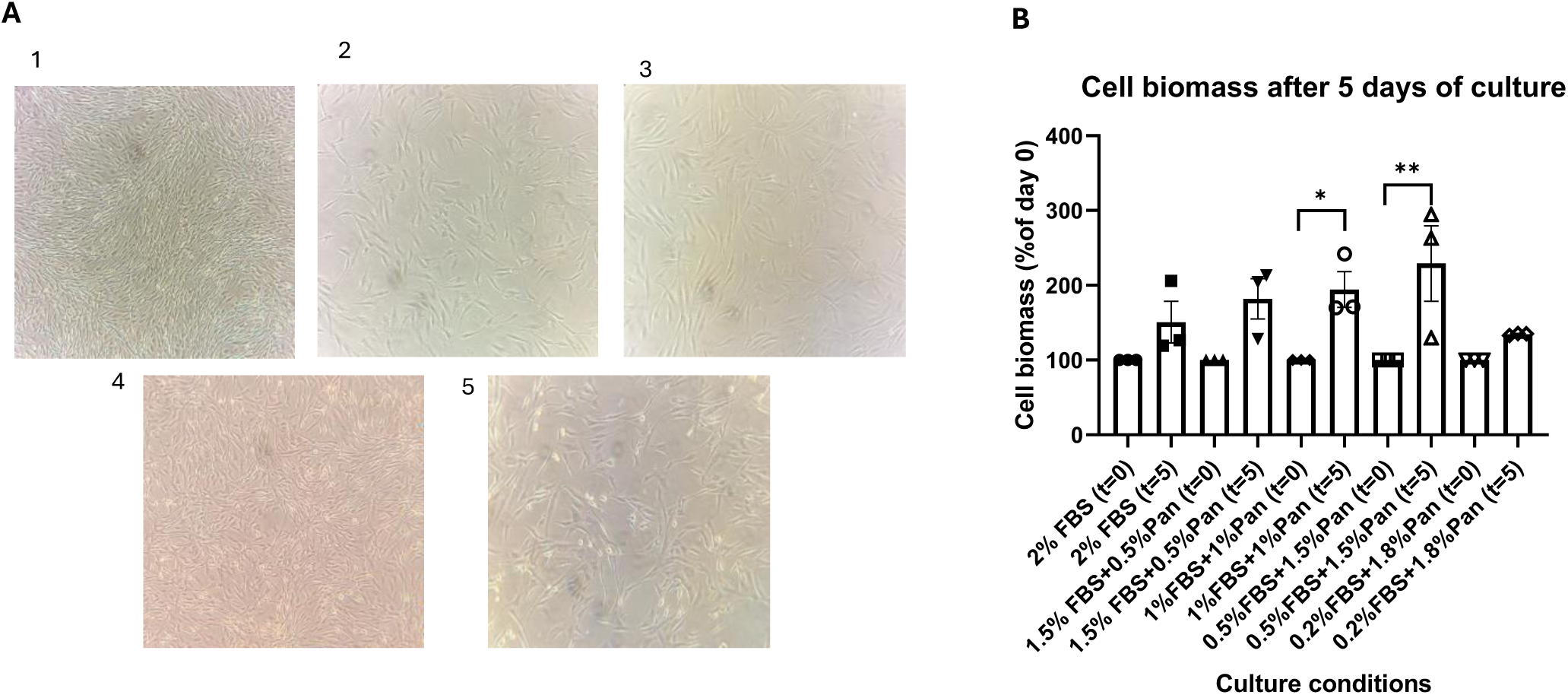
Optimization of Panexin supplementation for AT-MSC culture. (A) Representative phase-contrast images of AT-MSCs cultured under different Panexin/FBS supplementation conditions after 5 days of culture. Images correspond to the following culture media: (1) 2% FBS (in-house culture medium), (2) 1.5% FBS + 0.5% Panexin, (3) 1.0% FBS + 1.0% Panexin, (4) 0.5% FBS + 1.5% Panexin, and (5) 0.2% FBS + 1.8% Panexin. (B) Relative cell biomass determined by Crystal Violet staining at day 5. Data are presented as mean ± SEM (n = 3). *p < 0.05 and **p < 0.01 versus the corresponding day 0 control.

Under the standard culture condition (2% FBS), cell biomass increased after 5 days of culture, but no significant difference was observed. On the other hand, replacement of FBS with Panexin at proportions 1.0% Panexin and 1.0% FBS and 1.5% Panexin and 0.5% FBS, significantly increased cell biomass compared with their respective day 0 controls (p < 0.05 and p < 0.01, respectively). Among all tested conditions, 1.5% Panexin and 0.5% FBS yielded the highest cell biomass after 5 days of culture. In contrast, cultures supplemented with 1.8% Panexin and 0.2% FBS exhibited minimal cell growth over the 5-day culture period, with cell biomass remaining close to baseline levels. Representative images confirmed a markedly lower cell confluence than that observed under the other culture conditions, suggesting that this formulation was insufficient to sustain normal AT-MSC expansion. Based on these findings, the 1.5% Panexin and 0.5% FBS condition was selected for subsequent EV production experiments and attempting further reduction of FBS was discarded.

### 6. Serum BSA and EV identity marker detection on MSC-EVs derived from cells cultured in 1.5% Panexin + 0.5% FBS and 10% CDM-HD circulating media

Following the initial proteomic analyses and the assessment of albumin depletion during the transition from FBS-containing medium to CDM-HD, EV preparations obtained from subsequent hollow-fiber bioreactor experiments were further characterized by Western blot. EVs produced under the selected 1.5% Panexin + 0.5% FBS condition and under 10% CDM-HD were analyzed for albumin, the EV-associated markers CD81 and CD63, and the cellular marker calnexin. Lysed AT-MSCs were included as a positive control for calnexin detection, whereas FBS diluted 1:50 served as a positive control for BSA detection. No detectable albumin signal was observed in either EV preparation obtained under 1.5% Panexin + 0.5% FBS or 10% CDM-HD conditions (Figure 5A), whereas a clear signal was observed in the FBS control. These results are consistent with the absence of detectable FBS-derived albumin in the EV preparations obtained under the tested culture conditions. Both CD81 and CD63 were detected in EV preparations obtained under the two culture conditions (Figures 5B and 5C), with clearer signals observed than in the EV preparations analyzed during the previous FBS-to-CDM-HD transition. These results support the presence of EVs in both preparations. Calnexin was detected in the AT-MSC cell lysate control but was b arely detected in the EV preparations (Figure 5D). The absence of calnexin signal in the EV samples, together with the detection of CD81 and CD63, further supports the enrichment of EVs and the absence of detectable cellular protein contamination in the isolated preparations.

**Figure 5.**
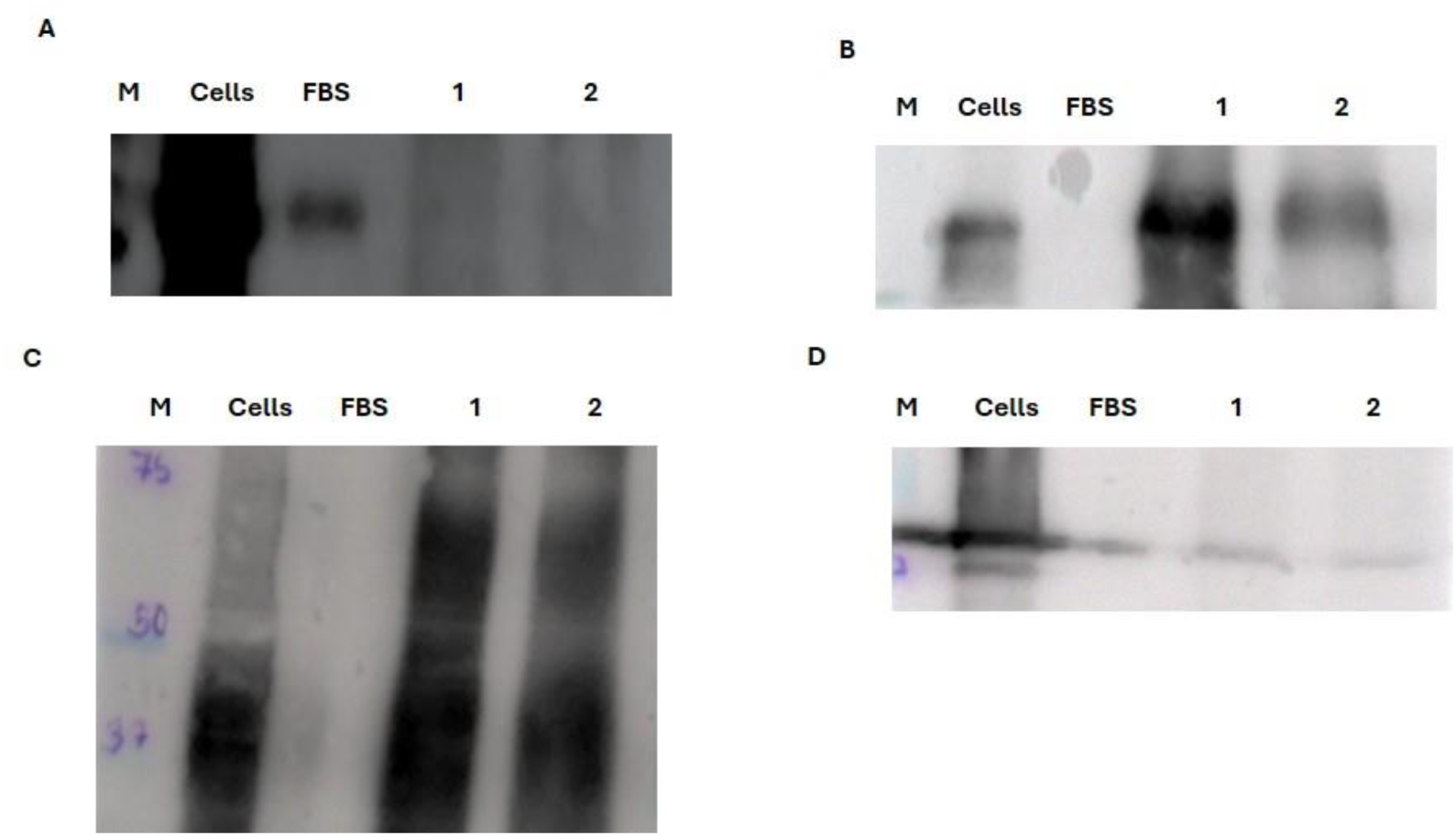
Western blot characterization of EVs produced under Panexin-and CDM-HD-based culture conditions. (A) Albumin immunoblot. (B) CD81 immunoblot. (C) CD63 immunoblot. (D) Calnexin immunoblot. Lane M, molecular weight marker; lane Cells, AT-MSC cell lysate; lane FBS, fetal bovine serum; lane 1, EVs produced under 1.5% Panexin + 0.5% FBS culture conditions; lane 2, EVs produced under 10% CDM-HD culture conditions.

### 7. Proteomic analysis of MSC-EVs derived from cells cultured in 1.5% Panexin + 0.5% FBS and 10% CDM-HD circulating media

Proteomic analysis made to AT-MSC-EVs isolated from cells cultured in 1.5% Panexin + 0.5% PBS identified a total of 1,551 protein groups. Based on organism annotation, 1,091 protein groups (70.3%) were exclusively assigned to *Homo sapiens*, 56 (3.6%) to *Bos taurus*, while 404 (26.0%) corresponded to protein groups containing both human and bovine assignments (Figure 6A). Another analysis was made to AT-MSC-EVs isolated from cells cultured in CDM-HD 10% identified a total of 1,044 protein groups. Based on organism annotation, 749 protein groups (71.2%) were exclusively assigned to *Homo sapiens*, 41 (3.9%) to *Bos taurus*, while 260 (24.9%) corresponded to protein groups containing both human and bovine assignments (Figure 6B). To characterize the most abundant proteins identified under each culture condition, the 35 highest-ranked protein groups were analyzed separately for CDM-HD and 1.5% Panexina + 0.5% FBS, with their corresponding protein annotations and EV-associated information summarized in Tables 4 and 5, respectively.

**Figure 6.**
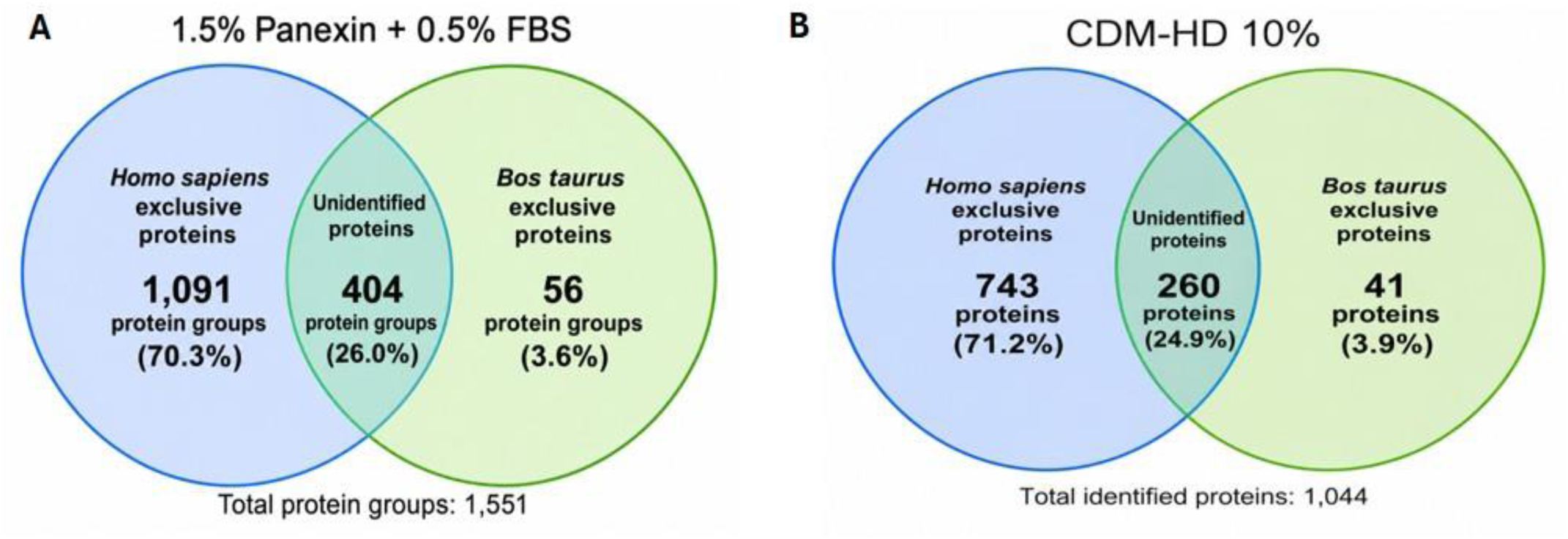
Comparative analysis of protein group distribution identified by proteomic analysis of extracellular vesicles (EVs) produced under different culture conditions. (A) EVs obtained in medium supplemented with 1.5% Panexin and 0.5% FBS, with 1,551 protein groups identified. (B) EVs obtained in medium supplemented with 10% CDM-HD, with 1,044 proteins identified. Venn diagrams show the distribution of protein groups classified as Homo sapiens-specific, Bos taurus-specific, or unidentified, expressed as the number of protein groups and the percentage of the total identified protein groups under each culture condition.

**Table 4.** Most abundant proteins identified in EVs produced under 1.5% Panexin + 0.5% FBS culture conditions. The 35 proteins with the highest abundance in the proteomic dataset are shown. Protein identity and organism assignment were based on UniProt annotations. EV relevance was classified according to reported evidence of association with EV preparations, extracellular matrix localization, or potential FBS-derived origin. Color coding: green, proteins unambiguously assigned to *Homo sapiens*; red, proteins unambiguously assigned to *Bos taurus*; yellow, proteins for which species assignment was ambiguous due to shared or non-specific peptide sequences between human and bovine proteins.

| **Ranking** | **UniProt ID** | **Protein name** | **Organism** | **EV relevance** |
|---|---|---|---|---|
| 1 | P13611 | Versican | *Homo sapiens* | EV-associated / ECM |
| 2 | Q08380 | Galectin-3-binding protein | *Homo sapiens* | EV-associated / extracellular |
| 3 | Q7Z7G0 | ABI family member 3-binding protein | *Homo sapiens* | EV-associated / ECM |
| 4 | P23142 | Fibulin-1 | *Homo sapiens* | EV-associated / ECM |
| 5 | G5E5A8 | Fibronectin (bovine) | *Bos taurus* | EV-associated / ECM |
| 6 | P02751 | Fibronectin | *Homo sapiens* | EV-associated / ECM |
| 7 | P11047 | Laminin subunit gamma-1 | *Homo sapiens* | EV-associated / ECM |
| 8 | P02452 | Collagen alpha-1(I) chain | *Homo sapiens* | EV-associated / ECM |
| 9 | P27487 | Dipeptidyl peptidase 4 (CD26) | *Homo sapiens* | EV-associated / membrane |
| 10 | P24821 | Tenascin-C | *Homo sapiens* | EV-associated / ECM |
| 11 | P04264 | Keratin 1 | *Homo sapiens* | EV-associated / cytoskeleton |
| 12 | P04439 | HLA class I histocompatibility antigen, A alpha chain | *Homo sapiens* | EV-associated / immune / MHC-I |
| 13 | P21589 | 5'-nucleotidase, ecto (CD73) | *Homo sapiens* | EV-associated / membrane / purinergic signaling |
| 14 | P08123 | Collagen alpha-2(I) chain | *Homo sapiens* | EV-associated / ECM |
| 15 | P07942 | Laminin subunit beta-1 | *Homo sapiens* | EV-associated / ECM |
| 16 | Q15113 | Procollagen C-endopeptidase enhancer 1 | *Homo sapiens* | EV-associated / ECM |
| 17 | P13645 | Keratin 10 | *Homo sapiens* | EV-associated / cytoskeleton |
| 18 | P98160 | Perlecan | *Homo sapiens* | EV-associated / ECM |
| 19 | P0CG47; P0CG48; P0CG53; P0CH28 | Polyubiquitin-B and Polyubiquitin-C | *Homo sapiens; Bos taurus* | EV-associated / ubiquitin system |
| 20 | Q16363 | Laminin subunit alpha-4 | *Homo sapiens* | EV-associated / ECM |
| 21 | O00468 | Agrin | *Homo sapiens* | EV-associated / ECM |
| 22 | P08962 | CD63 antigen | *Homo sapiens* | EV marker / membrane |
| 23 | P60709; P60712; P63258; P63261 | Cytoplasmic actin 1 and Cytoplasmic actin 2 | *Homo sapiens; Bos taurus* | EV-associated / cytoskeleton |
| 24 | P12111 | Collagen alpha-3(VI) chain | *Homo sapiens* | EV-associated / ECM |
| 25 | O00560 | Syntenin-1 | *Homo sapiens* | EV-associated / membrane / EV biogenesis |
| 26 | P15144 | Aminopeptidase N | *Homo sapiens* | EV-associated / membrane |
| 27 | A0A3Q1LKE5 | Tenascin-C (bovine) | *Bos taurus* | EV-associated / ECM |
| 28 | Q9Y6C2 | EMILIN-1 | *Homo sapiens* | EV-associated / ECM |
| 29 | P12109 | Collagen alpha-1(VI) chain | *Homo sapiens* | EV-associated / ECM |
| 30 | P12110 | Collagen alpha-2(VI) chain | *Homo sapiens* | EV-associated / ECM |
| 31 | P13646 | Keratin 13 | *Homo sapiens* | EV-associated / cytoskeleton |
| 32 | P02533 | Keratin 14 | *Homo sapiens* | EV-associated / cytoskeleton |
| 33 | P01889 | HLA class I histocompatibility antigen, B alpha chain | *Homo sapiens* | EV-associated / immune / MHC-I |
| 34 | Q12805 | Fibulin-3 | *Homo sapiens* | EV-associated / ECM |
| 35 | P07355 | Annexin A2 | *Homo sapiens* | EV-associated / membrane / cytoskeleton |

**Table 5.** Most abundant proteins identified in EVs produced under 10% CDM-HD culture conditions. The 35 proteins with the highest abundance in the proteomic dataset are shown. Protein identity and organism assignment were based on UniProt annotations. EV relevance was classified according to reported evidence of association with EV preparations, extracellular matrix localization, or potential FBS-derived origin. Color coding: green, proteins unambiguously assigned to *Homo sapiens*; red, proteins unambiguously assigned to *Bos taurus*; yellow, proteins for which species assignment was ambiguous due to shared or non-specific peptide sequences between human and bovine proteins.

| Ranking | UniProt ID | Protein name | Organism | EV relevance |
|---|---|---|---|---|
| 1 | Q08380 | Galectin-3-binding protein | *Homo sapiens* | EV-associated / extracellular |
| 2 | P13611 | Versican | *Homo sapiens* | EV-associated / ECM |
| 3 | P04264 | Keratin 1 | *Homo sapiens* | EV-associated / cytoskeleton |
| 4 | P13645 | Keratin 10 | *Homo sapiens* | EV-associated / cytoskeleton |
| 5 | P98160 | Perlecan | *Homo sapiens* | EV-associated / ECM |
| 6 | P02533 | Keratin 14 | *Homo sapiens* | EV-associated / cytoskeleton |
| 7 | P35527 | Keratin 9 | *Homo sapiens* | EV-associated / cytoskeleton |
| 8 | P35908 | Keratin 2 | *Homo sapiens* | EV-associated / cytoskeleton |
| 9 | Q7Z7G0 | ABI family member 3-binding protein | *Homo sapiens* | EV-associated / ECM |
| 10 | P08962 | CD63 antigen | *Homo sapiens* | EV marker / membrane |
| 11 | P27487 | Dipeptidyl peptidase 4 (CD26) | *Homo sapiens* | EV-associated / membrane |
| 12 | O00468 | Agrin | *Homo sapiens* | EV-associated / ECM |
| 13 | P11047 | Laminin subunit gamma-1 | *Homo sapiens* | EV-associated / ECM |
| 14 | P02538 | Keratin 6A | *Homo sapiens* | EV-associated / cytoskeleton |
| 15 | P23142 | Fibulin-1 | *Homo sapiens* | EV-associated / ECM |
| 16 | Q0P5J4 | Keratin 25 (bovine) | *Bos taurus* | EV-associated / cytoskeleton |
| 17 | P07585 | Decorin | *Homo sapiens* | EV-associated / ECM |
| 18 | P07942 | Laminin subunit beta-1 | *Homo sapiens* | EV-associated / ECM |
| 19 | Q16363 | Laminin subunit alpha-4 | *Homo sapiens* | EV-associated / ECM |
| 20 | P02792 | Ferritin light chain | *Homo sapiens* | EV-associated / iron homeostasis |
| 21 | P08670 | Vimentin | *Homo sapiens* | EV-associated / cytoskeleton |
| 22 | O00560 | Syntenin-1 | *Homo sapiens* | EV-associated / membrane / EV biogenesis |
| 23 | P13646 | Keratin 13 | *Homo sapiens* | EV-associated / cytoskeleton |
| 24 | P08473 | Neprilysin | *Homo sapiens* | EV-associated / membrane |
| 25 | P60709; P60712; P63258; P63261 | Cytoplasmic actin 1 and Cytoplasmic actin 2 | *Homo sapiens; Bos taurus* | EV-associated / cytoskeleton |
| 26 | P24821 | Tenascin-C | *Homo sapiens* | EV-associated / ECM |
| 27 | P34741 | Syndecan-2 | *Homo sapiens* | EV-associated / membrane / ECM |
| 28 | P15144 | Aminopeptidase N | *Homo sapiens* | EV-associated / membrane |
| 29 | P05556 | Integrin beta-1 | *Homo sapiens* | EV-associated / membrane / adhesion |
| 30 | Q14956 | Glycoprotein NMB | *Homo sapiens* | EV-associated / membrane |
| 31 | Q8N1N4 | Keratin 78 | *Homo sapiens* | EV-associated / cytoskeleton |
| 32 | Q14112 | Nidogen-2 | *Homo sapiens* | EV-associated / ECM |
| 33 | P81605 | Dermcidin | *Homo sapiens* | EV-associated / secreted / antimicrobial |
| 34 | P13647 | Keratin 5 | *Homo sapiens* | EV-associated / cytoskeleton |
| 35 | P0CG47; P0CG48; P0CG53; P0CH28 | Polyubiquitin-B and Polyubiquitin-C | *Homo sapiens; Bos taurus* | EV-associated / ubiquitin system |

**Table 6.** Comparison of the 35 most abundant proteins identified under 1.5% Panexin + 0.5% FBS and 10% CDM-HD culture conditions. The 35 proteins with the highest ranking under the 1.5% Panexin + 0.5% FBS condition were used as the reference set, and their corresponding ranking under 10% CDM-HD was determined. Rank change represents the difference between the rankings under the two conditions; positive values indicate an increase in relative ranking under 10% CDM-HD (↑), whereas negative values indicate a decrease (↓). Species assignment was based on UniProt annotation, with green indicating *Homo sapiens*, red indicating *Bos taurus*, and yellow indicating mixed human/bovine protein groups for which the protein group contains assignments to both species. Rankings represent relative positions within each individual proteomic dataset and should not be interpreted as direct comparisons of absolute protein abundance between conditions.

| **FBS rank** | **Protein name** | **UniProt ID** | **Organism** | **CDM-HD rank** | **Rank change** | **Movement** | **Interpretation** |
|---|---|---|---|---|---|---|---|
| 1 | Versican | P13611 | *Homo sapiens* | 2 | 1 | ↓ 1 | Moved down 1 positions |
| 2 | Galectin-3-binding protein | Q08380 | *Homo sapiens* | 1 | -1 | ↑ 1 | Moved up 1 positions |
| 3 | ABI family member 3-binding protein | Q7Z7G0 | *Homo sapiens* | 9 | 6 | ↓ 6 | Moved down 6 positions |
| 4 | Fibulin-1 | P23142 | *Homo sapiens* | 15 | 11 | ↓ 11 | Moved down 11 positions |
| 5 | Fibronectin (bovine) | G5E5A8 | *Bos taurus* | 64 | 59 | ↓ 59 | Moved down 59 positions |
| 6 | Fibronectin | P02751 | *Homo sapiens* | 80 | 74 | ↓ 74 | Moved down 74 positions |
| 7 | Laminin subunit gamma-1 | P11047 | *Homo sapiens* | 13 | 6 | ↓ 6 | Moved down 6 positions |
| 8 | Collagen alpha-1(I) chain | P02452 | *Homo sapiens* | 49 | 41 | ↓ 41 | Moved down 41 positions |
| 9 | Dipeptidyl peptidase 4 (CD26) | P27487 | *Homo sapiens* | 11 | 2 | ↓ 2 | Moved down 2 positions |
| 10 | Tenascin-C | P24821 | *Homo sapiens* | 26 | 16 | ↓ 16 | Moved down 16 positions |
| 11 | Keratin 1 | P04264 | *Homo sapiens* | 3 | -8 | ↑ 8 | Moved up 8 positions |
| 12 | HLA class I histocompatibility antigen, A alpha chain | P04439 | *Homo sapiens* | 55 | 43 | ↓ 43 | Moved down 43 positions |
| 13 | 5'-nucleotidase, ecto (CD73) | P21589 | *Homo sapiens* | 42 | 29 | ↓ 29 | Moved down 29 positions |
| 14 | Collagen alpha-2(I) chain | P08123 | *Homo sapiens* | 69 | 55 | ↓ 55 | Moved down 55 positions |
| 15 | Laminin subunit beta-1 | P07942 | *Homo sapiens* | 18 | 3 | ↓ 3 | Moved down 3 positions |
| 16 | Procollagen C-endopeptidase enhancer 1 | Q15113 | *Homo sapiens* | 59 | 43 | ↓ 43 | Moved down 43 positions |
| 17 | Keratin 10 | P13645 | *Homo sapiens* | 4 | -13 | ↑ 13 | Moved up 13 positions |
| 18 | Perlecan | P98160 | *Homo sapiens* | 5 | -13 | ↑ 13 | Moved up 13 positions |
| 19 | Polyubiquitin-B and Polyubiquitin-C | P0CG47; P0CG48; P0CG53; P0CH28 | *Homo sapiens; Bos taurus* | 35 | 16 | ↓ 16 | Moved down 16 positions |
| 20 | Laminin subunit alpha-4 | Q16363 | *Homo sapiens* | 19 | -1 | ↑ 1 | Moved up 1 positions |
| 21 | Agrin | O00468 | *Homo sapiens* | 12 | -9 | ↑ 9 | Moved up 9 positions |
| 22 | CD63 antigen | P08962 | *Homo sapiens* | 10 | -12 | ↑ 12 | Moved up 12 positions |
| 23 | Cytoplasmic actin 1 and Cytoplasmic actin 2 | P60709; P60712; P63258; P63261 | *Homo sapiens; Bos taurus* | 25 | 2 | ↓ 2 | Moved down 2 positions |
| 24 | Collagen alpha-3(VI) chain | P12111 | *Homo sapiens* | 51 | 27 | ↓ 27 | Moved down 27 positions |
| 25 | Syntenin-1 | O00560 | *Homo sapiens* | 22 | -3 | ↑ 3 | Moved up 3 positions |
| 26 | Aminopeptidase N | P15144 | *Homo sapiens* | 28 | 2 | ↓ 2 | Moved down 2 positions |
| 27 | Tenascin-C (bovine) | A0A3Q1LKE5 | *Bos taurus* | 41 | 14 | ↓ 14 | Moved down 14 positions |
| 28 | EMILIN-1 | Q9Y6C2 | *Homo sapiens* | 54 | 26 | ↓ 26 | Moved down 26 positions |
| 29 | Collagen alpha-1(VI) chain | P12109 | *Homo sapiens* | 61 | 32 | ↓ 32 | Moved down 32 positions |
| 30 | Collagen alpha-2(VI) chain | P12110 | *Homo sapiens* | 99 | 69 | ↓ 69 | Moved down 69 positions |
| 31 | Keratin 13 | P13646 | *Homo sapiens* | 23 | -8 | ↑ 8 | Moved up 8 positions |
| 32 | Keratin 14 | P02533 | *Homo sapiens* | 6 | -26 | ↑ 26 | Moved up 26 positions |
| 33 | HLA class I histocompatibility antigen, B alpha chain | P01889 | *Homo sapiens* | 122 | 89 | ↓ 89 | Moved down 89 positions |
| 34 | Fibulin-3 | Q12805 | *Homo sapiens* | 193 | 159 | ↓ 159 | Moved down 159 positions |
| 35 | Annexin A2 | P07355 | *Homo sapiens* | 40 | 5 | ↓ 5 | Moved down 5 positions |

The Top 35 proteins identified under CDM-HD were predominantly of human origin, comprising 32 human, one bovine, and two mixed human/bovine protein groups (Table 4). The highest-ranked proteins included galectin-3-binding protein and versican, together with several keratins and extracellular matrix (ECM)-associated proteins. Among the most relevant EV-and ECM-associated proteins identified in this ranking were CD63, perlecan, laminin subunits, agrin, and syntenin-1. Overall, the ranking was strongly dominated by proteins of human origin.

A similarly human-dominated profile was observed for the 1.5% Panexin + 0.5% FBS condition, with 31 human, two bovine, and two mixed human/bovine protein groups among the Top 35 (Table 5). Versican and galectin-3-binding protein remained among the most abundant proteins, while the ranking also contained several ECM-associated proteins, including fibulin-1, laminins, collagens, and tenascin-C. Notably, both human and bovine fibronectin were among the most highly ranked proteins in this condition. Since fibronectin was not used as an adhesion substrate in this bioreactor system, the detection of bovine fibronectin cannot be attributed to fibronectin used during cell attachment and is therefore particularly relevant when considering the contribution of serum-derived proteins.

To evaluate changes in the relative abundance of the most highly ranked proteins between conditions, the 35 proteins ranked highest under 1.5% Panexina + 0.5% FBS were tracked across the complete proteomic ranking obtained under CDM-HD. Several proteins maintained a similar high ranking between conditions. Versican and galectin-3-binding protein, for example, occupied the first two positions under 1.5% Panexina + 0.5% FBS and remained at positions 2 and 1, respectively, under CDM-HD(Table. 6).

In contrast, several ECM-associated proteins showed substantial decreases in ranking under CDM-HD. Bovine fibronectin moved from rank 5 to rank 64, while human fibronectin moved from rank 6 to rank 80. Collagen alpha-1(I) chain and collagen alpha-2(I) chain similarly shifted from ranks 8 and 14 to ranks 49 and 69, respectively. Fibulin-3 showed one of the largest changes, moving from rank 34 to rank 193. These changes indicate that several ECM-associated proteins that were highly represented under the Panexina/FBS condition occupied substantially lower positions under CDM-HD.

Conversely, several cytoskeletal and EV-associated proteins increased their relative ranking under CDM-HD. Keratin 1, keratin 10, and keratin 14 moved from ranks 11, 17, and 32 to ranks 3, 4, and 6, respectively. Perlecan and CD63 also showed marked increases in ranking, moving from positions 18 and 22 to 5 and 10, respectively. This shift suggests a different distribution of the most abundant EV-associated proteins between the two culture conditions.

The comparison of bovine proteins revealed an additional notable pattern. Bovine fibronectin and bovine tenascin-C were both highly ranked under 1.5% Panexina + 0.5% FBS but occupied substantially lower positions under CDM-HD. In contrast, BSA showed the opposite behavior, moving from rank 161 under 1.5% Panexina + 0.5% FBS to rank 63 under CDM-HD. This was unexpected given the presence of FBS in the former condition and highlights that the abundance of individual serum-derived proteins did not necessarily follow the expected pattern based solely on the presence or absence of FBS.

Human albumin also showed a marked change in ranking, moving from rank 1,061 under 1.5% Panexina + 0.5% FBS to rank 101 under CDM-HD. Together, these observations highlight substantial differences in the relative representation of individual proteins between the two culture conditions, including proteins of both human and bovine origin.

### 8. ELISA protein quantification shows evidence of BSA cross-contamination in the BE-SEC column

To evaluate residual BSA contamination in AT-MSC-EV preparations, albumin levels were quantified by ELISA in samples obtained under different culture conditions and at different stages of the EV production and purification process. Initially, EV preparations obtained using DMEM supplemented with 1.5% Panexin and 0.5% FBS were compared with those produced under chemically defined medium (CDM) conditions. Both preparations contained detectable levels of BSA, with mean values of approximately 0.75 ng and 0.90 ng per EV preparation, respectively. However, no statistically significant difference was observed between the two culture conditions (Figure 7A).

**Figure 7.**
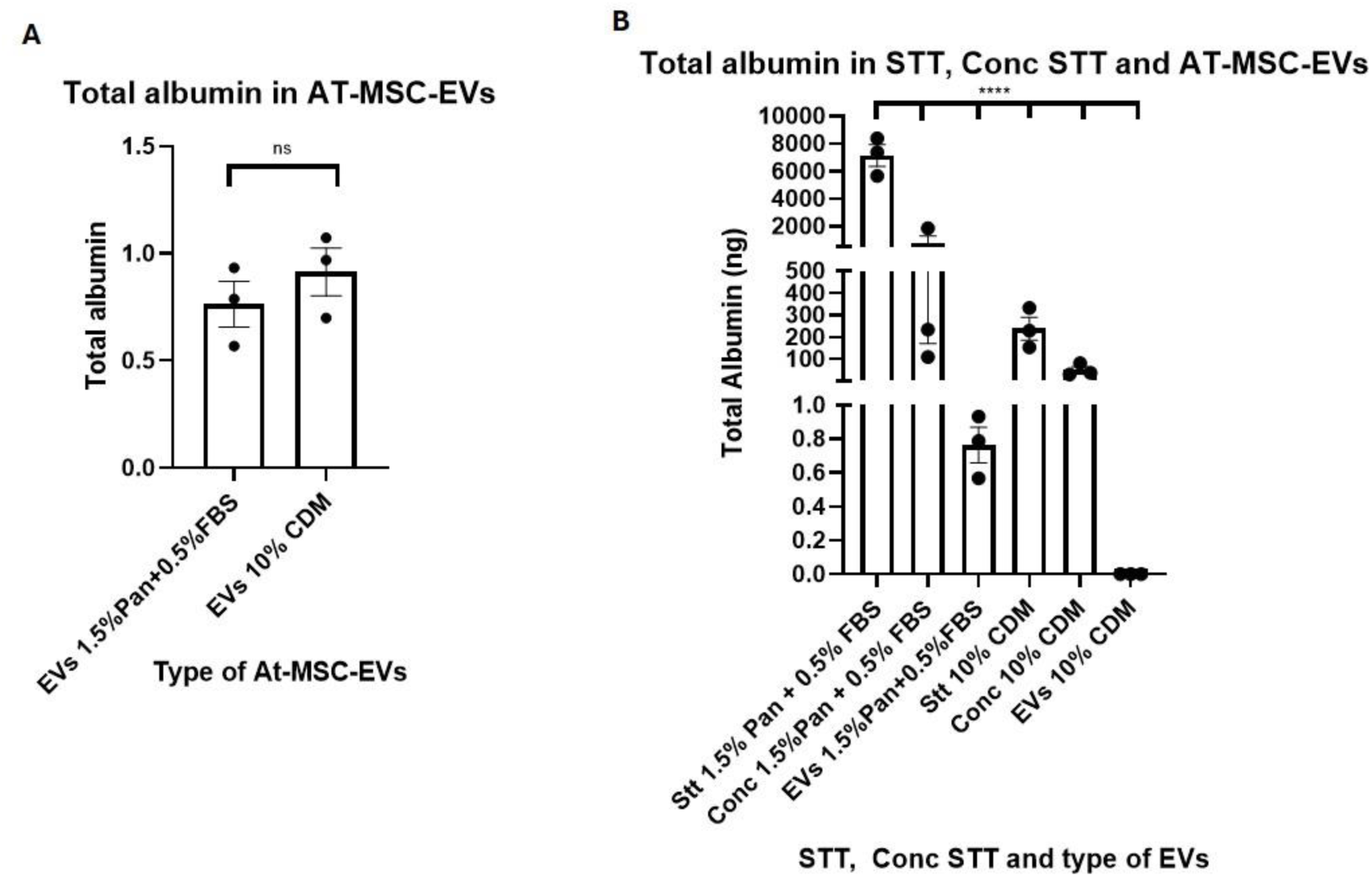
Quantification of residual BSA in AT-MSC-EV preparations and samples collected during the purification process. (A) Total BSA (ng per EV preparation) measured by ELISA in EVs produced under two different culture conditions: 1.5% Panexin + 0.5% FBS and 10% CDM. No statistically significant differences were observed between conditions. (B) Total BSA measured by ELISA in conditioned supernatants (STT) before and after tangential flow filtration (TFF) concentration (Conc STT), and in the final AT-MSC-EV preparations obtained under 1.5% Panexin + 0.5% FBS and 10% CDM-HD culture conditions. The initial supernatant volume (40 mL) was concentrated to approximately 5 mL before FPLC purification. Albumin levels progressively decreased throughout the purification process. In the final CDM-HD-derived EV preparations, albumin levels were below the lower limit of quantification of the assay (0.69 ng/mL). Data are presented as mean ± SEM. ****p < 0.0001.

To further investigate the reduction of albumin during EV processing, BSA levels were quantified in the initial conditioned supernatants (STT), after tangential flow filtration (TFF) concentration (Conc STT), and in the final AT-MSC-EV preparations obtained under both culture conditions (Figure 7B).

The highest total albumin levels were detected in the initial supernatant obtained from cultures supplemented with 1.5% Panexin and 0.5% FBS. Following TFF concentration, a marked reduction in the total amount of recovered albumin was observed. A similar trend was observed under CDM-HD conditions, although the initial albumin levels were considerably lower. Albumin levels decreased further following FPLC purification, with residual albumin remaining detectable in the final EV preparation obtained under 1.5% Panexin and 0.5% FBS conditions. In contrast, albumin levels in the initial CDM-HD-derived EV preparation were below the lower limit of quantification of the assay (0.69 ng/mL). Statistical analysis revealed highly significant differences between the compared sample groups (****, p < 0.0001; Figure 7B).

To investigate the residual albumin initially detected in the CDM-HD-derived EV preparation, the possibility of contamination associated with the chromatography column was considered, as FBS-containing samples had previously been processed using the same column. Therefore, the purification process was repeated using a new Capto Core 700 column, and three independent CDM-HD-derived EV preparations were analyzed by ELISA. In all three preparations, BSA levels were below the lower limit of quantification of the assay (0.69 ng/mL). These findings suggest that the albumin initially detected in the CDM-HD-derived EV preparation may have resulted from residual carryover from the previously used chromatography column rather than from the CDM-HD culture conditions themselves.

## DISCUSSION

The use of fetal bovine serum (FBS) during the expansion and production of cell-derived extracellular vesicles (EVs) represents an important consideration when assessing their protein composition, as serum-derived components may contribute to the proteins detected in the final EV preparation. To investigate the extent of this contribution under conventional serum-containing conditions, EVs were produced from AT-MSCs cultured in a hollow-fiber bioreactor cartridge with a 20 KDa cut-off membrane in the presence of 10% FBS, expecting a minimal bovine protein carryover, and subsequently subjected to an initial proteomic analysis. On the contrary, the analysis revealed a substantial contribution of bovine-derived proteins, with the most abundant serum protein, bovine serum albumin (66 KDa), ranking among the most abundant proteins identified. The presence of bovine proteins in human cell culture-derived samples has previously been associated with FBS supplementation, and their contribution may complicate the interpretation of proteomic datasets, particularly because homologous human and bovine proteins can display high sequence similarity (Abramowicz et al. 2018; Sugár et al. 2020). Indeed, bovine serum proteins have been reported as abundant contaminants in human cell culture-derived samples and may even be incorrectly assigned to human proteins when species-specific sequences cannot be unequivocally distinguished (Sugár et al. 2020).

To further investigate the contribution of serum-derived proteins suggested by the proteomic analysis, BSA was subsequently evaluated by Western blot during the transition from FBS-containing medium to CDM-HD. The detection of albumin under FBS-containing conditions, followed by its reduction and eventual absence after adaptation to CDM-HD, provided an independent confirmation of the serum-derived protein contribution observed in the proteomic analysis. The use of BSA as an indicator of serum-derived protein carryover is particularly relevant in EV studies, as bovine serum albumin has been identified as a major potential contaminant in EV preparations and can contribute to detection errors during downstream mass spectrometry analyses (Abramowicz et al. 2018). Thus, the concordance between the proteomic and Western blot analyses supports the contribution of FBS-derived proteins to the protein profile of EV preparations generated under serum-containing conditions.

The presence of abundant serum-derived proteins may have consequences beyond their detection as contaminants, as they can interfere with the characterization of less abundant EV-associated proteins. Previous studies have shown that highly abundant serum proteins, particularly albumin, can limit the detection of lower-abundance proteins in untargeted proteomic analyses and reduce the coverage of the EV proteome (Abramowicz et al. 2018; Smolarz et al. 2019; Reymond et al. 2023). Similarly, the low abundance of EV-associated proteins relative to abundant serum proteins such as albumin and apolipoproteins has been reported as a major challenge for EV proteomic characterization(Lane et al. 2019). In this context, the reduction in albumin observed by Western blot during adaptation to CDM-HD was accompanied by a clearer detection of the EV-associated markers CD81 and CD63, suggesting that reducing the contribution of abundant serum proteins may facilitate the detection of EV-associated proteins. Furthermore, the increased number of proteins identified by proteomic analysis under CDM-HD conditions compared with FBS-containing conditions provides additional evidence that reducing serum-derived protein contamination may improve proteomic coverage. This is consistent with previous observations showing that serum-derived proteins can mask lower-abundance exosome-specific components and compromise the representation of the EV proteome in mass spectrometry analyses (Abramowicz et al. 2018).

The transition to CDM-HD resulted in a markedly different protein profile compared with the EV preparations generated under 10% FBS conditions. The strong reduction in the contribution of bovine proteins, together with the substantial decrease in the relative abundance of albumin, indicates that replacing the serum-containing medium effectively minimized the contribution of serum-derived proteins to the isolated end product. This observation is consistent with previous studies showing that the use of serum-free or chemically defined culture conditions can reduce the contribution of medium-derived components to MSC-EV preparations and result in a more clearly defined EV-associated protein profile (Pachler et al. 2017; Hanai et al. 2023). In addition, differences in the culture medium have been shown to influence the molecular characteristics of MSC-derived EVs, indicating that changes in the culture environment can extend beyond the removal of exogenous serum proteins (Zhang et al. 2025).

Despite the marked reduction in serum-derived proteins following the transition to CDM-HD, a subset of bovine proteins remained detectable after prolonged culture under the defined-medium condition. This persistence may not necessarily reflect incomplete removal of FBS from the culture medium. Previous studies have shown that serum proteins can adsorb to surfaces exposed to fetal calf serum and remain associated with those surfaces after their initial exposure (Haas and Culp 1982). Furthermore, the composition of the adsorbed protein layer can differ from that of the original serum, indicating that individual serum proteins may interact differently with the surface (Hannan and Reilly 1988). Therefore, surface-associated serum proteins may represent a potential source of residual bovine proteins detected after subsequent medium replacement and washing steps.

This possibility may be particularly relevant to the present bioreactor system, in which the hollow fibers were coated with human fibronectin before cell seeding, as recommended by the manufacturer. Fibronectin is known to interact with several plasma proteins, and immobilized fibronectin can retain soluble proteins through specific protein–protein interactions. For example, immobilized fibronectin has been shown to bind the complement protein C1q, while plasma gelsolin has also been shown to interact with immobilized fibronectin (Bing et al., n.d.; Lind and Janmey 1984). In addition, Hannan and Reilly demonstrated that proteins adsorbed from fetal serum to surfaces can participate in secondary interactions involving fibronectin, suggesting that the protein layer formed on a serum-exposed surface may be more complex than the proteins initially adsorbed directly to the surface (Hannan and Reilly 1988). Based on these observations, we hypothesize that exposure of the fibronectin-coated hollow fibers to FBS during the initial culture period may have promoted the association and retention of specific bovine serum proteins on the fiber surface. Among the bovine proteins that remained detectable after adaptation to CDM-HD were albumin, alpha-2-HS-glycoprotein, alpha-1-antiproteinase, tenascin C, and E3 ubiquitin-protein ligase FANCL. These proteins could subsequently remain associated with the bioreactor despite medium replacement and washing steps, providing a possible explanation for the residual detection of selected bovine proteins after adaptation to CDM-HD.

Importantly, the interaction of serum proteins with fibronectin is not expected to be uniform across different proteins. Vaidya and Ofoli showed that fibronectin adsorbs rapidly and essentially irreversibly at an interface, whereas albumin exhibited more reversible adsorption and could be displaced by fibronectin. These differences indicate that the persistence of individual serum proteins on a fibronectin-coated surface may depend on their specific adsorption and interaction properties (Vaidya and Ofoli 2005). This may be relevant to our findings, as albumin showed a marked reduction following adaptation to CDM-HD, whereas a subset of bovine proteins remained detectable. Although the retention of these proteins by the fibronectin coating was not directly assessed in the present study, their persistence may therefore be compatible with differential interactions between serum proteins and the fibronectin-coated bioreactor surface.

The culture medium represents an important upstream factor in extracellular vesicle (EV) production, as its composition can influence the molecular characteristics of the resulting EV preparations. In this context, chemically defined media (CDM) offer an alternative to serum-containing formulations by providing culture conditions without the need for fetal bovine serum (FBS). This is particularly relevant for EV production, as serum-derived proteins may remain as contaminants in the final EV preparations. In addition, components derived from culture medium supplements may co-isolate with EVs and introduce exogenous molecules into the final preparations, potentially interfering with their characterization (Auber et al. 2019). Different chemically defined media formulations have been shown to affect the EV proteome and the abundance of EV-associated surface markers, indicating that changes in medium composition can modify EV characteristics beyond the simple presence or absence of serum (Zhu et al. 2021). In addition, components derived from culture medium supplements may co-purify with EV preparations and interfere with the characterization of EV-associated molecules, highlighting the importance of considering the contribution of exogenous components when interpreting EV analyses (Auber et al. 2019b). These considerations are particularly relevant from a manufacturing perspective, as avoiding the introduction of bovine serum proteins into the final EV product is important for the development of GMP-compatible production processes. Therefore, following the insufficient reduction in serum-derived proteins observed after adaptation to CDM-HD, an alternative culture strategy combining 1.5% Panexin with 0.5% FBS was evaluated aiming to further reduce the contribution of serum-derived proteins to the EV-associated protein profile. It is important to note that CDM-HD is a chemically defined media only for high-density 3D cultures that can not be used in the initial steps of the hollow fiber bioreactor setup.

Because reducing FBS exposure may affect cell growth and consequently the production of cell-derived EVs, different Panexin/FBS ratios attempted during the adaptation procedure were first evaluated in AT-MSC cultures to identify a condition that allowed continued cell growth while reducing the proportion of FBS to the minimum. In our experiments, both 1.0% Panexin + 1.0% FBS and 1.5% Panexin + 0.5% FBS supported a significant increase in cell biomass over the 5-day culture period, whereas the 1.8% Panexin + 0.2% FBS condition showed minimal cell growth. The 1.5% Panexin + 0.5% FBS condition produced the highest cell biomass among the conditions evaluated, even superior to FBS 2%, and was therefore selected for subsequent MSC bioreactor culture. This approach allowed us to reduce the FBS concentration while maintaining AT-MSC growth under the conditions tested.

The absence of detectable albumin by Western blot analysis in EV preparations generated under both 1.5% Panexin + 0.5% FBS and CDM-HD conditions indicates a substantial reduction in serum-derived albumin compared with the FBS-containing condition. In both preparations, CD81 and CD63 were detected, whereas the cellular marker calnexin was barely detected, supporting the presence of EV-enriched preparations without cellular protein contamination. These findings indicate that the reduction in FBS associated with the Panexin-based condition was sufficient to eliminate detectable albumin by Western blot while enhacing detection of EV-associated markers.

Despite the absence of detectable BSA by Western blot, the proteomic analysis of EV preparations generated with 1.5% Panexin + 0.5% FBS revealed a residual contribution of bovine proteins. The predominance of human proteins among the most highly ranked proteins indicates that the reduction in FBS substantially decreased the representation of serum-derived proteins; however, the detection of bovine proteins, including bovine fibronectin, demonstrates that this contribution was not completely eliminated. This finding is consistent with previous reports showing that reducing or removing serum-derived components can decrease, but not necessarily completely prevent, the contribution of bovine material to human cell-derived EV preparations (Pham et al. 2021b). Importantly, the detection of bovine proteins by proteomics despite the absence of an albumin signal by Western blot highlights the higher sensitivity of proteomic analyses and that the absence of detectable albumin by immunoblot does not necessarily indicate the complete absence of bovine-derived proteins in the EV preparation.

The comparison between the 1.5% Panexin + 0.5% FBS and 10% CDM-HD conditions further revealed differences in the relative representation of individual proteins, despite both conditions producing predominantly human protein profiles. Several extracellular matrix-associated proteins that were highly ranked under the Panexin/FBS condition showed substantially lower rankings under CDM-HD, whereas other proteins, including several cytoskeletal and EV-associated proteins, increased in relative ranking under CDM-HD. These differences suggest that the effect of changing the culture medium extends beyond the reduction of serum-derived proteins and may influence the relative protein composition of the resulting EV preparations. This interpretation is consistent with previous studies demonstrating that culture medium formulation can modify the proteomic composition and molecular characteristics of EVs (Zhu et al. 2021b). However, because protein rankings represent relative positions within each individual proteomic dataset, these differences should not be interpreted as direct evidence of changes in absolute protein abundance between the two conditions.

To further investigate the residual BSA detected in the proteomic analyses, its presence was quantified by a specific BSA ELISA in the final EV preparations and at different stages of the purification process. Although Western blot analysis did not reveal detectable albumin in preparations obtained using either 1.5% Panexin + 0.5% FBS or CDM-HD, ELISA detected low amounts of BSA in both conditions, with no significant differences between them. This finding suggests that residual albumin was present at levels below the detection capability of the immunoblot under the conditions used. The detection of albumin in EV preparations is relevant because free proteins may co-isolate with EVs during purification and interfere with the assessment of preparation purity (Stolk et al. 2015; Ter-Ovanesyan et al., 2023.). However, proteins such as albumin may also associate with the EV surface as part of the protein corona, rather than simply representing co-isolated contaminants(Tóth et al. 2021). This protein layer may influence EV interactions with recipient cells, cellular uptake and in vivo distribution, as demonstrated for MSC-derived EVs (Liam-Or et al. 2024). Therefore, reducing the incorporation of exogenous bovine proteins while considering the contribution of human-derived proteins to the EV-associated protein corona may be relevant for preserving the biological properties of EV preparations. Moreover, protein-derived contaminants from chemically defined collection media may also contribute to the protein content of isolated EV fractions, indicating that the use of serum-free conditions does not necessarily prevent protein co-isolation (Phelps et al. 2026).

Analysis of the conditioned supernatants before and after TFF showed a marked reduction in the total amount of albumin recovered following concentration in both culture conditions, despite the reduction in sample volume from approximately 40 to 5 mL. This suggests that a substantial proportion of the BSA present in the initial supernatant was eliminated by the TFF procedure, despite of the cartridge 15 KDa cut-off. Since BSA was not quantified in an intermediate sample collected after FPLC, the contribution of each purification step to its overall reduction cannot be determined. Nevertheless, the low BSA levels detected in the final EV preparations are consistent with further removal of free proteins during downstream purification. In this regard, Ter-Ovanesyan et al. (2023) demonstrated that Capto Core 700, when incorporated into a multimodal chromatography column, can contribute to the removal of free proteins, including albumin, from EV preparations. Although their study used plasma-derived EVs and a combined chromatographic system, it provides a rationale for the potential contribution of this resin to protein removal in our purification workflow.

An unexpected finding was the detection of albumin in the initial CDM-derived EV preparation, despite the absence of FBS during the collection phase. Since the Capto Core 700 column had previously been used to process FBS-containing samples, residual albumin retained in the column was considered a possible source of carryover. To investigate this possibility, the purification was repeated using a new column. In all three independent CDM-derived EV preparations, albumin levels were below the lower limit of quantification of the ELISA assay (0.69 ng/mL). These results support the possibility that the albumin initially detected under CDM conditions resulted from residual carryover associated with the previously used column rather than from the culture medium itself. However, as residual albumin was not directly measured on the original column, this explanation remains a plausible interpretation rather than definitive evidence of the contamination source.

Overall, this study demonstrates the optimization of an AT-MSC-EV production and isolation workflow aimed at obtaining high-purity EV preparations with minimal bovine protein contamination. This was achieved by combining a chemically defined medium during the initial cell expansion phase with CDM-HD during the final production phase, followed by a purification strategy involving TFF, FPLC using a BE-SEC approach, and ultrafiltration. The resulting EV preparations showed no detectable BSA by ELISA, supporting the effectiveness of this approach in reducing serum-derived protein contamination. Taken together, these findings provide a basis for further developing a scalable EV production process compatible with GMP manufacturing requirements.

## ACKNOWLEDGEMENTS

The authors would like to thank Estela Herranz for her valuable technical assistance and support throughout this study. Funding: *Industrial PhD Grant, Comunidad Autónoma de Madrid IND2023/BMD-27323* and *Ministerio de Ciencia, e Innovación y Universidades y Fondo Europeo de Desarrollo Regional PID2021-126274OB-I00; CNS2022-135368*. Alan Rivera-Tenorio was supported by *Industrial PhD Grant, Comunidad Autónoma de Madrid IND2023/BMD-27323.* Miriam M. Rodríguez de Lope was supported by *Ayudas para la realización del Programa Investigo, Comunidad Autónoma de Madrid (INV/09-PIN1-00013.4/2022)*.

